# Molecular basis for the activation of the Fatty Acid Kinase complex of *Staphylococcus aureus*

**DOI:** 10.1101/2024.03.19.585040

**Authors:** Megan J. Myers, Zhen Xu, Benjamin J. Ryan, Zachary R. DeMars, Miranda J. Ridder, David K. Johnson, Christina N. Krute, Tony S. Flynn, Maithri M. Kashipathy, Kevin P. Battaile, Nicholas Schnicker, Scott Lovell, Bret D. Freudenthal, Jeffrey L. Bose

**Affiliations:** Department of Microbiology, Molecular Genetics and Immunology, University of Kansas Medical Center, Kansas City, Kansas, USA; Protein and Crystallography Facility, University of Iowa, Iowa City, Iowa, USA; Department of Biochemistry and Molecular Biology, University of Kansas Medical Center, Kansas City, Kansas, USA; Computational Chemical Biology Core, University of Kansas, Lawrence, Kansas, USA; Protein Structure & X-Ray Crystallography Laboratory, University of Kansas, Lawrence, Kansas, USA; NYX, New York Structural Biology Center, Upton, NY 10027, USA

**Keywords:** MRSA, *S. aureus*, proline isomerization, fatty acid metabolism, fatty acid kinase

## Abstract

Gram-positive bacteria utilize a Fatty Acid Kinase (FAK) complex to harvest fatty acids from the environment. The complex, consisting of the fatty acid kinase, FakA, and an acyl carrier protein, FakB, is known to impact virulence and disease outcomes. However, FAK’s structure and enzymatic mechanism remain poorly understood. Here, we used a combination of modeling, biochemical, and cell-based approaches to establish critical details of FAK activity. Solved structures of the apo and ligand-bound FakA kinase domain captured the protein state through ATP hydrolysis. Additionally, targeted mutagenesis of an understudied FakA Middle domain identified critical residues within a metal-binding pocket that contribute to FakA dimer stability and protein function. Regarding the complex, we demonstrated nanomolar affinity between FakA and FakB and generated computational models of the complex’s quaternary structure. Together, these data provide critical insight into the structure and function of the FAK complex which is essential for understanding its mechanism.

## Introduction

*Staphylococcus aureus* is a Gram-positive bacterium asymptomatically colonizing around 30% of the population ^1^. In addition, it is the leading cause of skin infections and can cause a myriad of diseases with high morbidity and mortality in both the nosocomial and community environments ^2, 3, 4, 5^. Indeed, it is a global health problem estimated to kill one million people worldwide annually ^6^. This is due to the plethora of virulence factors produced by the bacterium, its metabolic diversity to adapt to various niches, and the high frequency of antibiotic resistance ^7, 8, 9, 10, 11, 12, 13, 14, 15, 16^. New options to combat *S. aureus* infections are critically needed, which requires a more detailed understanding of the physiology, pathogenesis, and virulence of this pathogen. Due to differences between bacterial and host pathways, fatty acid metabolism and biosynthesis are attractive targets for novel therapeutic development.

Fatty acids are an essential component of life as they are the building blocks of lipid membranes. They are common to the bacterial environment during infection as part of host cells, as a component of low-density lipoproteins, and as free fatty acids. This is particularly relevant to skin and soft tissue infections as host saturated and unsaturated fatty acids (UFAs) are ample, with the latter known to be antimicrobial ^17, 18, 19^. Because of the abundance of fatty acids within the host environment, bacteria have adopted ways to acquire exogenous fatty acids (exoFAs). ExoFAs can serve to supplement bacterial endogenous fatty acid biosynthesis by the FASII system or be degraded by β-oxidation. Because of this, exoFA acquisition contributes to resistance to FASII inhibitors, such as Triclosan ^20, 21, 22, 23^. As discovered in *S. aureus*, Gram-positive bacteria can acquire exoFAs using the Fatty Acid Kinase (FAK) complex which imports and activates exoFAs to be primarily inserted into lipids ^8, 9^. The FAK complex consists of FakA and FakB where the former is an ATP-utilizing kinase that phosphorylates the head group of fatty acids brought in by the acyl carrier protein, FakB ^24^. *S. aureus* contains two FakB proteins. FakB1 prefers saturated fatty acids and has been proposed to scavenge saturated exoFAs or recycle those made or released by *S. aureus* ^25, 26^. FakB2 preferentially interacts with UFAs and thus binds host fatty acids since *S. aureus* does not synthesize UFAs ^26, 27^. In this process, exoFAs passively diffuse into the bacterial membrane and are bound by FakB to be brought into the cytosol ^25^. Here, FakB interacts with FakA to form the FAK complex, whereby the kinase activity of FakA phosphorylates the fatty acid bound to FakB. This is processed by the phospholipid synthesis protein PlsY to insert the fatty acid primarily into the *sn1* stereospecific position of phospholipids ^28^. Without the FAK complex, *S. aureus* is unable to utilize exoFAs to synthesize lipids ^27, 28^.

FakA was originally discovered due to its impact on *S. aureus* virulence factor production and was termed VfrB; since then, it was determined to be a fatty acid kinase and renamed FakA ^8^. The FAK complex is important for a variety of *S. aureus* cellular processes including susceptibility to host fatty acids, membrane fluidity, toxin production, and virulence depending on the host niche ^8, 9, 10, 11, 12, 13, 14, 15, 16^. Despite these studies demonstrating the importance of FAK in *S. aureus*, much is still not known about the functional complex. Indeed, the basic understanding of the composition of the complex and the mechanism by which fatty acid phosphorylation occurs is unknown. Previous structural studies identified key aspects of the FakA and FakB proteins but have not captured how the complex forms the functional unit in response to binding the respective ligands, ATP and FA, and undergoes catalysis ^29, 30^.

Here, we used a combination of biochemical, structural, and cellular approaches to elucidate the mechanism of the functional FAK complex in *S. aureus*. Importantly, this study identifies the molecular changes in FakA upon binding ATP that forms the active site. Additionally, we identified the importance of a Zn-binding pocket in the maintenance of the FakA homodimeric state and the formation of the functional FAK complex. Together these results provide crucial missing information about the activity of FakA, structural changes that occur within the FakA kinase domain to allow the assembly of a functional complex, and the impact of these elements on FAK complex activity.

## Results

### FakA Domain Architecture

To better understand the mechanism of FakA we employed X-ray crystallography. However, efforts toward the full-length structure were unsuccessful. We thus used a combination of limited proteolysis and AlphaFold modeling to predict domains of FakA to interrogate individually ^31^. FakA was determined to possess three domains separated by unstructured regions (Fig. 1A and B). We believe the unstructured regions contributed to our inability to solve the full structure using crystallography. Therefore, we truncated the protein into individual domains to obtain structures of the FakA C- and N-terminal domains. The N-terminal domain of FakA, termed the kinase domain, spanning from residue 1 to 210 was crystallized in several states to provide mechanistic insight during the catalytic cycle of FakA. These include the apo form (Apo-FakA_N), the ATP analog bound form (AMP-PNP-FakA_N), and the ADP bound form (ADP-FakA_N). The organization of the FakA N-terminal domain in our structures agrees with recently solved structures of the ligand-bound FakA N-term ^29, 30^. Each of these structures formed a similar overall fold comprising a bundle of eight α-helices termed α1: G7-L31, α2: T41-N58, α3: I64-G78, α4: G81-I98, α5: S106-K122, α6: I132-N149, α7: C153-N173 and α9: S186-L202 and a short helix termed α8: A176-V181 (Fig. 2A and B). The kinase active site sits at the top (as orientated in Fig. 2A) of this helical barrel near the short helix α8.

**Figure 1.**
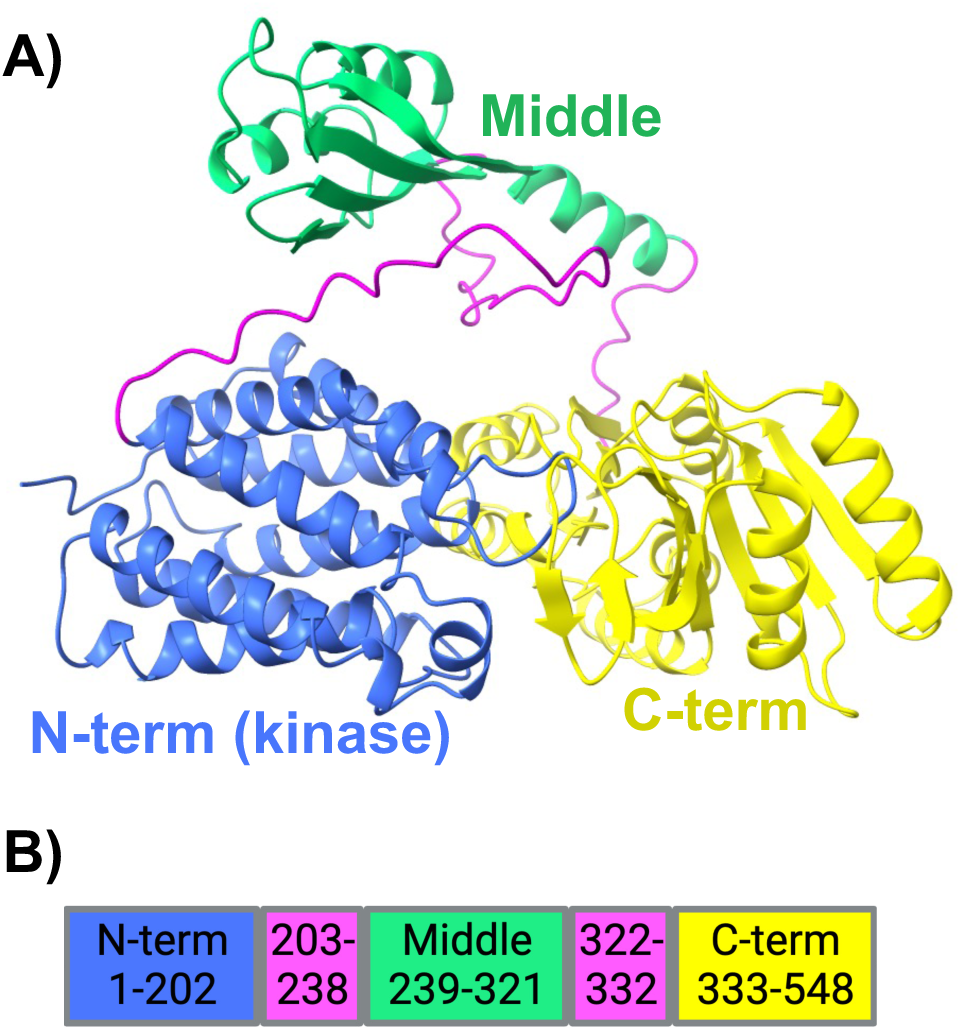
FakA monomer structure. **A)** FakA AlphaFold predicted model. Blue: N-terminal kinase domain, Green: Middle domain, Yellow: C-terminal domain, Magenta: Unstructured regions. **B)** Predicted amino acid residues in each domain or interdomain region.

**Figure 2:**
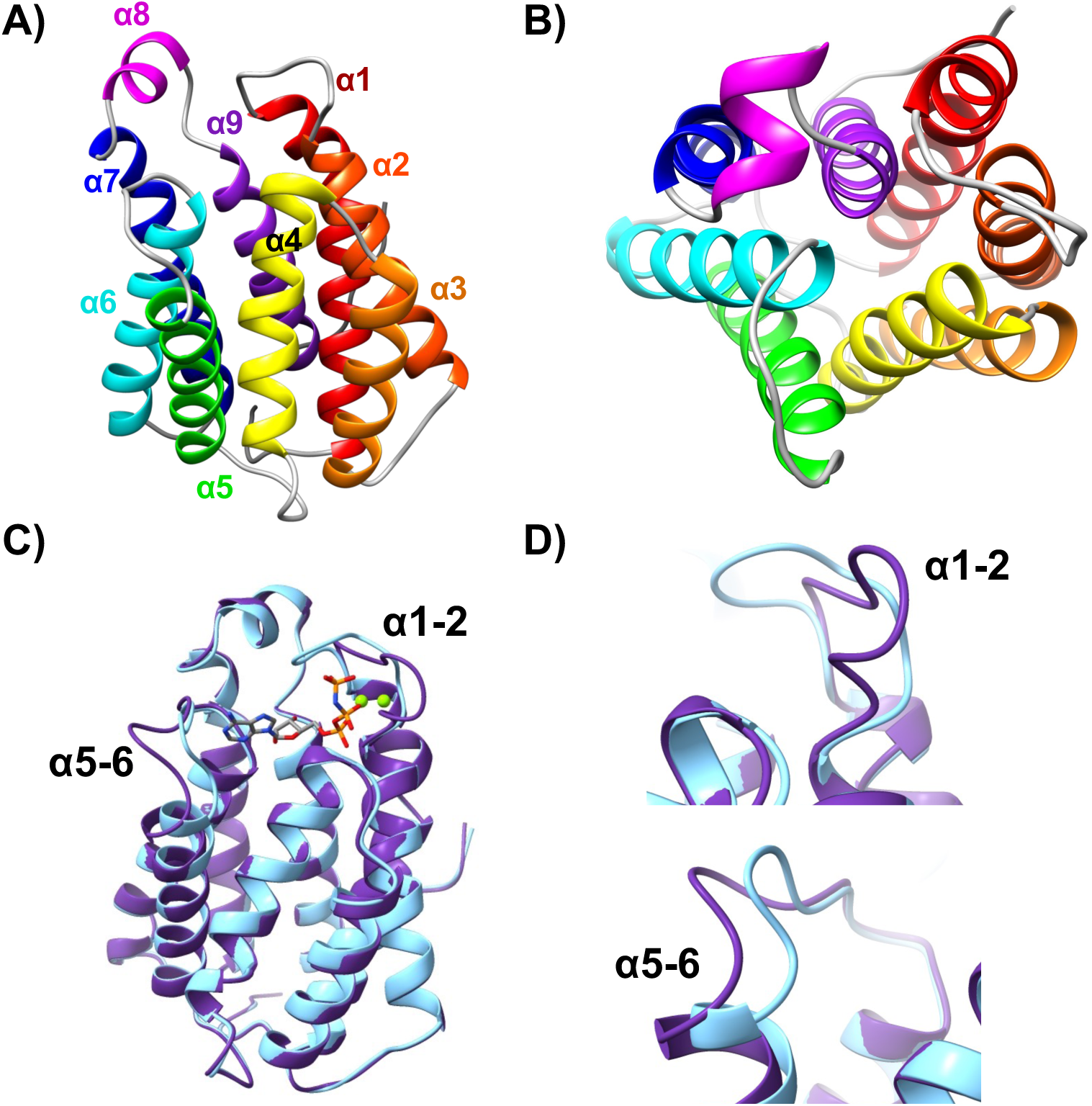
N-terminal FakA crystal structure. **A)** Apo-FakA crystal structure with α-helices colored from N-term to C-term from red to violet. **B)** 90° rotation relative to panel A to view down α-helical barrel. **C)** Overlay of Apo-FakA_N (purple) and AMP-PNP-FakA_N (light blue) with AMP-PNP labeled in gray and colored by element. Mg^2+^ ions are rendered as green spheres. Loops connecting α-helices 1 and 2 as well as 5 and 6 are labeled. **D)** Zoomed in view of the α1-2 (top) and α5-6 (bottom) loop movement from apo to bound.

To understand the changes that occur upon ligand binding to FakA, we solved the structure of Apo-FakA_N to 1.90 Å resolution and compared it to the ligand-bound AMP-PNP-FakA_N solved to 1.15 Å resolution (Fig. 2 and Movie S1). Superposition of Apo-and AMP-PNP-FakA_N yielded an RMSD deviation of 0.48 Å between Ca atoms (201 residues) indicating relatively little difference in the overall structure of the α-helical barrel (Fig. 2C). However, major differences were observed in the loops connecting α1/α2 and α5/α6 (Fig. 2D). Upon ATP binding, the α5-6 loop moves toward the ligand binding pocket to stabilize the nucleotide. The α1/2 loop extends away from the ATP binding pocket in the apo form, then collapses to bring key residues in proximity to the γ-phosphate of ATP and to accommodate FakB binding.

By comparing the Apo-FakA_N to AMP-PNP-FakA_N, we observed distinct localized structural changes upon ATP binding. These changes stabilize the nucleobase, sugar, and triphosphate portion of ATP in the active state as well as prepare a hydrophobic area for binding to FakB. One of these changes is a significant movement of the loop between α1-2, we term an activating loop, consisting of residues Leu31-Thr41 (Fig. 2C and D). Of note, Pro35 shifts 10 Å upon ATP binding and isomerizes from the *trans* to *cis* form. In this conformation, Pro35 forms a stabilizing interaction with the aromatic ring of Tyr34, closing over the adenosine base of the bound ATP to form the top of the ATP binding pocket in FakA (Fig. 3A-C, and Movie S2). The loop between α5/α6 is shifted by ∼5 Å upon binding AMP-PNP (Fig. 2C and D) to form hydrogen bonds between the backbone carbonyl of Lys126 to the N6 position, the backbone of Val128 to the N1 position, and Thr131 side chain to the N3 position of the nucleobase (Fig. 3A-C, S1A, and Movie S2). This, along with a side chain conformational change and hydrogen bond in Asn82, forms the nucleobase binding pocket providing specificity for adenosine (Fig. 3A-C, S1A, and Movie S2). All other residues in this α5/α6 loop region are similar in both structures. Additionally, hydrogen bonds between the sugar portion of the nucleotide and FakA_N are made between the backbone of Ile132 and O2, the side group of Asp185 with O2 and O3, and the side chain of Ser186 and O3 on the nucleotide (Fig. 3A-C, S1A, and Movie S2). At the triphosphate portion of the nucleotide, the shift of the α1-2 loop creates several stabilizing hydrogen bonds through the following contacts: the backbone nitrogen of Val36 to oxygen (O1G) on the γ-phosphate, between the sidechain of Thr41 to α-phosphate oxygen-1 (O1A), the sidechain of Asn44 to the α-phosphate oxygen-2 (O2A), the side chain and backbone of Ser83 to O2A, and the backbone and sidechain of Asn82 hydrogen bonds to O1B and N3B, respectfully (Fig. 3D-F, S1B, and Movie S3). Upon ATP binding, Asp38 and Asp40 within the α1-2 loop reposition 8.25 Å and 2.81 Å, respectively to coordinate the two Mg^2+^ ions (Fig. 3G-I and Movie S4). The two Mg^2+^ coordinate the α-, β-, γ-phosphate of the AMP-PNP as well as the catalytic triad Asp38, Asp40, and Asn32 (Fig. 3G-I and Movie S4). To verify these amino acids are involved in kinase activity, we used a malachite green Kinase assay. No activity was observed when FakA, FakB2, the fatty acid oleic acid, or ATP were omitted (Fig. 4A). In agreement with Asp38 and Asp40 being part of the catalytic site, no activity was measured when both Asp residues were substituted with Ala (Fig. 4B), consistent with our prior *in vivo* work showing these residues are critical for FakA function ^10^.

**Figure 3:**
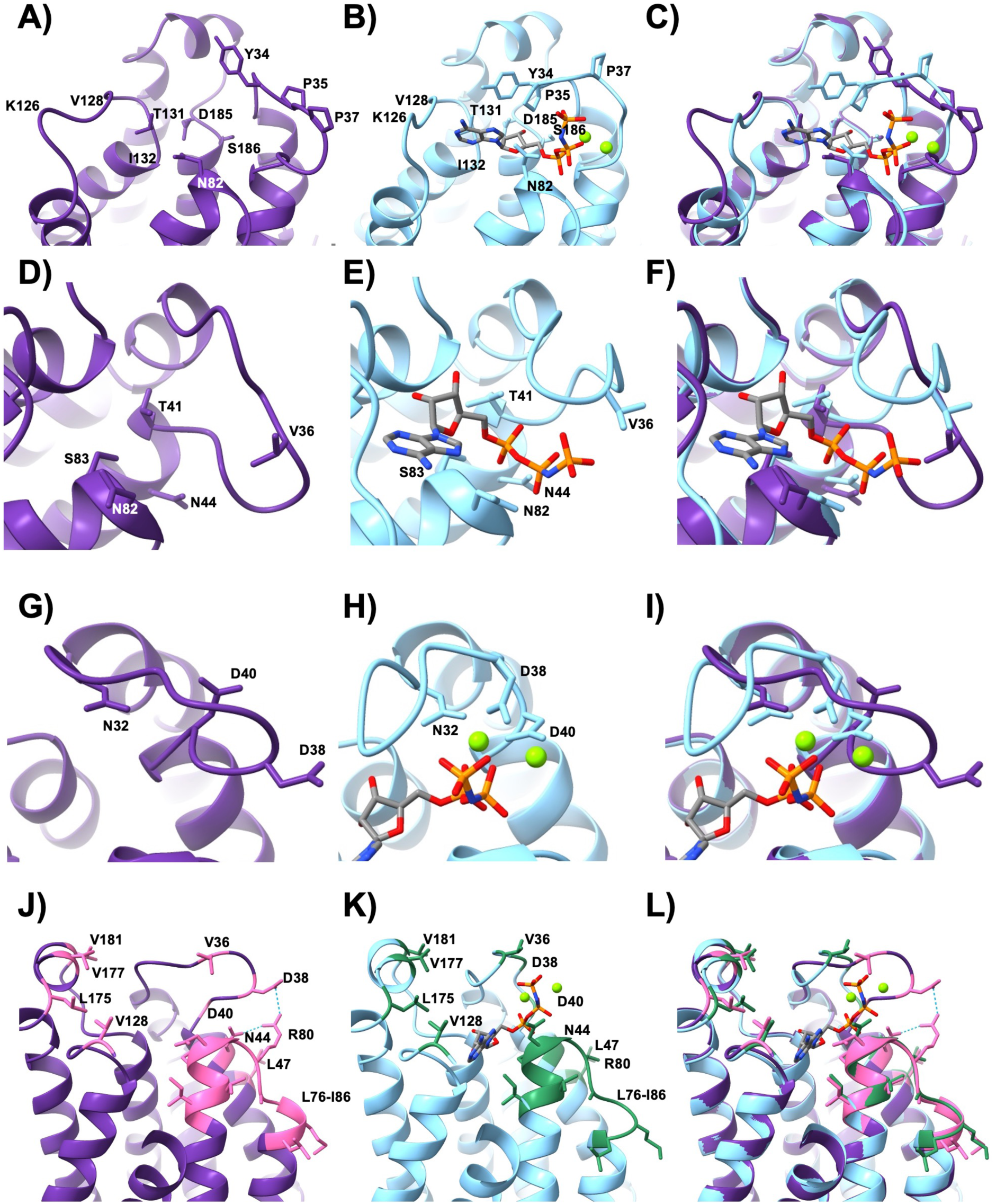
N-terminal FakA regions of interest. **A-C)** Nucleobase and sugar-binding pocket of FakA_N. **D-F)** Triphosphate binding of FakA_N. **G-H)** Mg^2+^ coordination by FakA_N. **J-L)** Altered binding face of FakA_N for FakB interaction. Apo-FakA_N is rendered in purple, and AMP-PNP-FakA_N is rendered in blue. Mg^2+^ ions are rendered as green spheres. AMP-PNP is rendered in gray cylinders with important atoms colored by element. In J-L, conserved hydrophobic residues are rendered in pink or green for Apo- and AMP-PNP-FakA_N, respectively.

**Figure 4:**
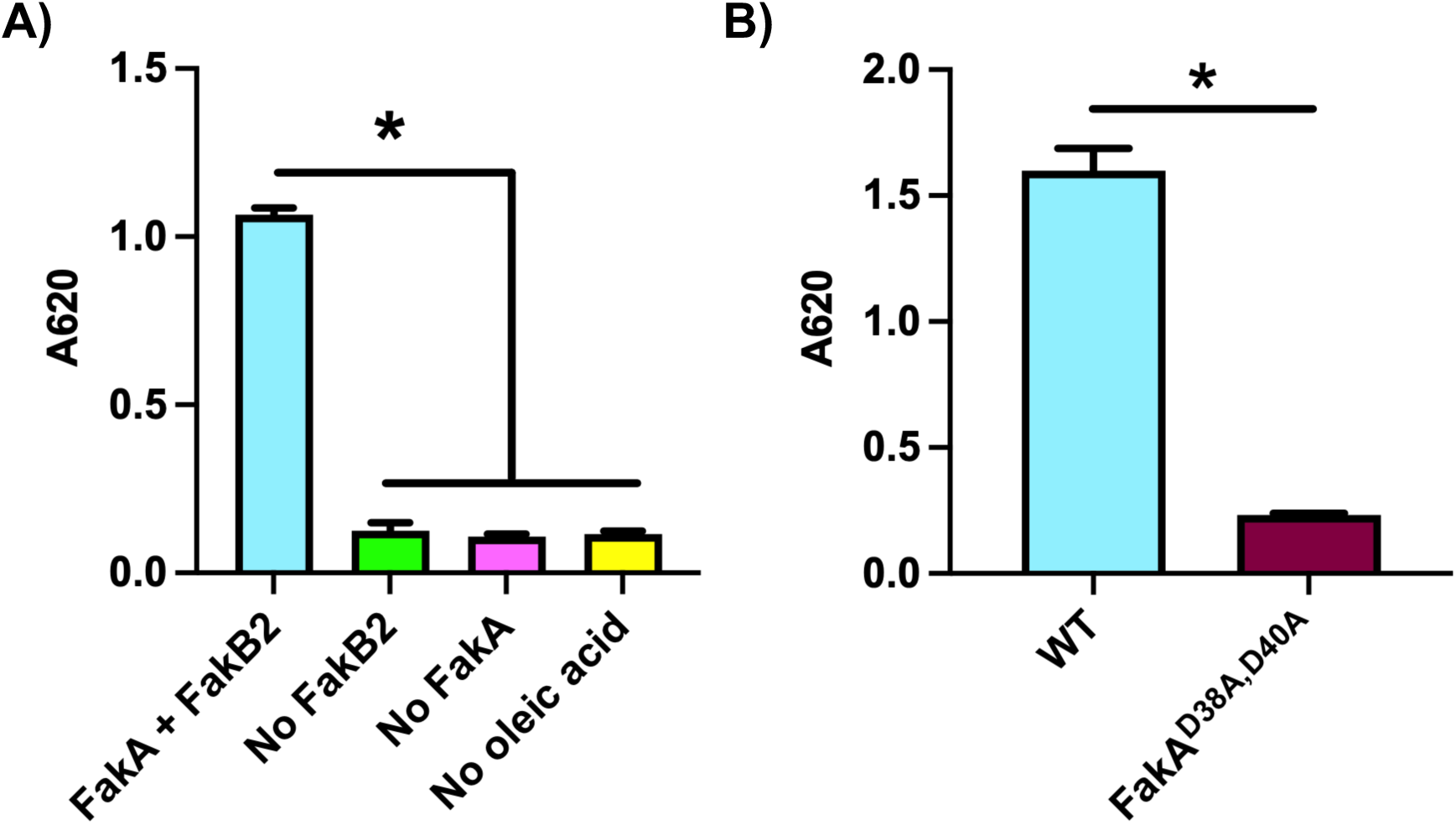
Asp38 and Asp40 are critical for FakA kinase activity. **A)** Kinase assay for the FAK complex activity or controls of no FakB, no FakA, or no oleic acid. **B)** Kinase assay showing activity is a loss in FakA^D28A, D40A^ variant in which Asp38 and Asp40 were replaced with Alanine. All proteins are His-tagged. Bars representative of the mean (n=2 for (A) or 4 for (B)) with SEM from a representative experiment. * indicates p<0.01 by Welch’s t-test.

Comparing the Apo- and AMP-PNP-FakA_N structures also gives insight into the structural changes that would accommodate FakB binding. Upon ATP binding, the repositioning of Val36 from the interior of the protein outward after the shift in the α1-2 loop creates a new binding face that we expect FakB to recognize and helps orient the fatty acid substrate in proximity to the γ-phosphate of ATP (Fig. 3J-L, S2, and Movie S5). The movement of this activating loop exposes a group of conserved hydrophobic residues including Val181, Val177, Leu175, Val128, Leu47, and the loop between Leu76-Ile86 when FakA is bound to ATP, ready to bind FakB (Fig. 3J-L, S2, and Movie S5). In Apo-FakA_N, the α1-2 loop protrudes away from the FakA ligand binding site; we predict this prevents premature binding of FakB (Fig. 3J-L). Hydrogen bonds between Asn44 and Asp38 with Arg80 stabilize this open form of FakA N-term, before ATP binding, then collapse and destabilize when in the ATP bound, closed form (Fig. 3J-L and Movie S5). Our interpretation is that the open state prevents FakB from binding to FakA that is not bound by ATP.

To determine the post-catalysis product state of the kinase catalytic cycle, we utilized ADP as a ligand to determine the ADP-metal-bound FakA structure to 1.25 Å resolution (Fig. S3). The structure of ADP-FakA_N was nearly identical to AMP-PNP-FakA_N aside from the absence of the γ-phosphate as superposition yielded an RMSD of 0.178 Å (Fig. S3). ADP is likely then released to return FakA to the apo state after the fatty acid is phosphorylated. Our crystallography data provides snapshots of the structural changes that occur in the FakA active site following the binding of ATP during the kinase catalytic cycle.

Having solved the N-terminal domain, we turned to the C-terminal FakA domain. We purified, crystallized, and solved the structure of the truncated C-terminal FakA protein (amino acids 328-548) to 1.55 Å resolution (Fig. S4). This matches the published C-term FakA structure (PDB 6W6B, ^30^). The FakA C-term structure consists of both β- sheets and α-helices (Fig. S4). In our initial discovery of FakA (originally called VfrB), we modeled this domain to have high structural similarity to a *Thermatoga* fatty acid binding protein. Additionally, C-terminal FakA is structurally similar to its acyl-binding protein partner, FakB. The solved crystal structures of the N- and C-terminal FakA domains overlay with RMSDs of 0.485 Å and 0.753 Å, respectively, to our original AlphaFold model, further corroborating our results (Fig. S5). To assess the fatty acid binding capability of the FakA C-terminal domain, we attempted adding fatty acids to the crystallization conditions but were unable to obtain a bound structure. Based on our structure and others ^30^, while the C-terminal domain of FakA resembles a fatty acid-binding protein, it appears unable to bind fatty acids alone.

### Middle domain of FakA

The Middle domain of FakA is predicted to contain α-helices and β-sheets (Fig. 5A). Conservation analysis of the region revealed three highly conserved amino acids within this domain: His282, His284, and Cys240 (Fig. S6). Further examination of the AlphaFold predicted structure of these amino acids shows their orientation into a pocket that we hypothesized would bind a metal to be a functional scaffold supporting the structure of the Middle domain and the biological function of full-length FakA (Fig. 5A). To investigate the presence of a metal ion, we compared purified FakA to a buffer control. The greatest observed difference between the buffer control and protein samples was a 55-fold increase in the amount of zinc present in FakA (Fig. 5B). To rule out that the His-tag of the purified FakA may be binding the zinc, rather than a metal binding pocket, we purified an untagged FakA and repeated the ICP-MS. Consistent with the previous result, this purified FakA also contained zinc (Fig. 5C). To isolate binding to the Middle domain, we purified the C-terminal FakA with and without the Middle domain and found that only the truncated protein containing the Middle domain had Zn (Fig. 5C). Finally, to distinguish if the putative metal-binding pocket identified in the AlphaFold model was responsible for Zn binding, we purified a FakA^H282A, H284A^ mutant and a FakA^C240A^ mutant. Mutation of C240 led to a significant decrease in the amount of zinc and this was further decreased in the FakA^H282A, H284A^ mutant (Fig. 5C).

**Figure 5:**
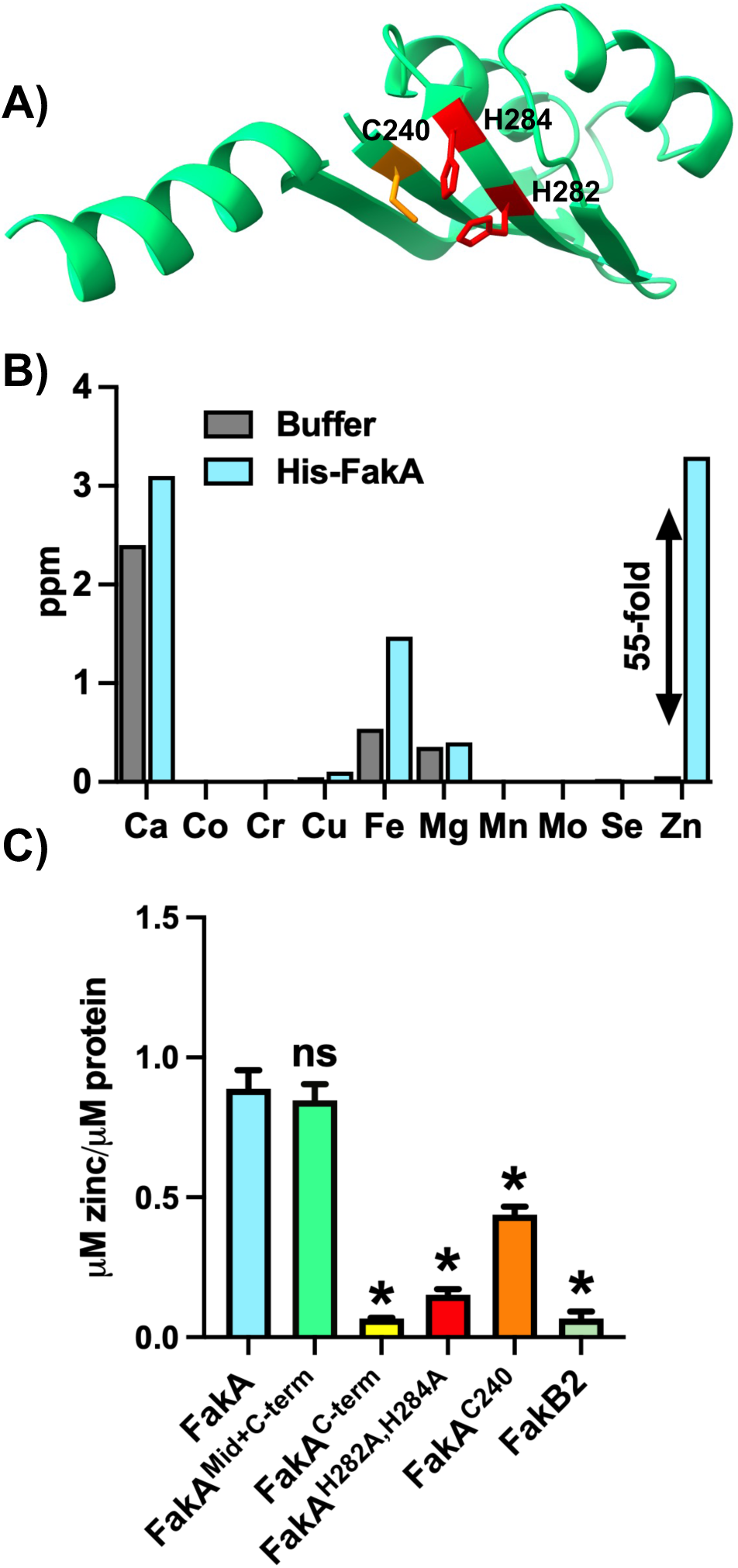
FakA Middle domain binds zinc. **A)** AlphaFold model of Middle domain with C240, H282, and H284 shown in red. **B)** ICP-MS metal quantification in parts per million (ppm) analysis of purified His-tagged FakA and buffer control. **C)** ICP-MS analysis of zinc associated with the indicated proteins. Untagged FakA is used in panel C. * indicates p<0.01 by one-way ANOVA compared to full-length wild-type FakA.

To test whether Zn binding impacted FakA kinase activity, we performed a kinase assay on the purified proteins. The FakA^C240A^ and FakA^H282A, H284A^ substitutions each led to a loss of FakA kinase activity (Fig. 6A). Next, we wanted to test if our biochemical loss of FakA kinase activity translated to a biological difference in FakA activity *in vivo* using two assays previously shown to be impacted by FakA activity: α-toxin production and linoleic acid resistance ^8, 12^. As previously shown, the *fakA* mutant showed a lack of α-toxin activity compared to the wild-type strain. When either C240 or both H282 and H284 were replaced with Ala, hemolysis activity was not restored, demonstrating that these amino acids are critical to FakA activity (Fig. 6B). We then tested whether FakA^C240A^ and FakA^H282A, H284A^ would restore the sensitivity of the *fakA* mutant to linoleic acid. The linoleic acid resistance of these variants was similar to that of the *fakA* mutant (Fig. 6C-D, and S7). Together, these data demonstrate that FakA possesses a metal binding pocket in the Middle domain that is essential to FakA kinase activity *in vitro* and FakA function *in vivo*. We predict that disruption of this metal binding pocket disrupts the integrity of the Middle domain, rendering the protein non-functional.

**Figure 6:**
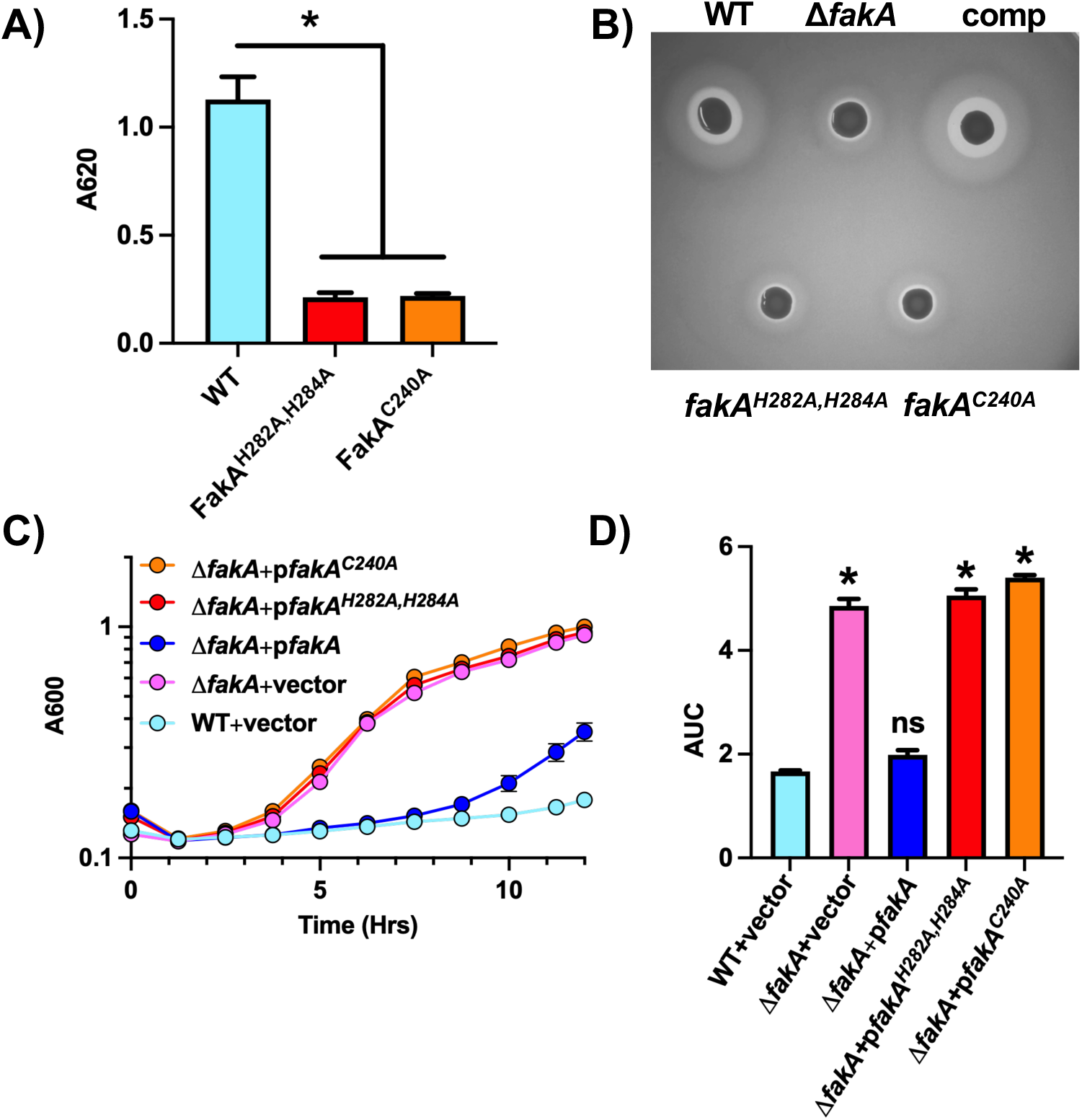
Metal coordination within Middle domain is essential for FakA function. **A)** FAK complex kinase assay indicating loss of kinase activity in variants in which Cys^240^ was replaced with Ala and when His^282^ and His^284^ were replaced with Ala. All proteins were purified using a GST tag which was removed prior to assay. Bars represent the mean (n=4) with SEM. **B)** Rabbit blood hemolysis (clearing around colony) of WT, *fakA* mutant, or mutant with a plasmid expressing WT FakA (comp), FakA^H282A, H284A^, or FakA^C240A^. **C)** Growth of variants in TSB supplemented with 192 µM (0.006%) linoleic acid. Symbols represent that mean (n=3) with SEM. **D)** Area under the curve analysis of panel C. Data are from a representative experiment. * indicates p-value <0.01 by Welch’s t-test.

### FakA Homodimerization

Because many kinases function as dimers, we next wanted to investigate FakA’s functional oligomeric state and the potential regulatory implications. In support of this, we previously showed by analytical ultracentrifugation that FakA purified as a dimer ^9^, though the structural basis of the dimerization was not known. AlphaFold-Multimer ^31^ modeling indicates dimerization is likely to occur through specific amino acid interactions in the Middle domain (Fig. 7A). To experimentally examine the FakA dimer structure, we employed several approaches. First, mass photometry of purified FakA reveals that 77% of the protein was within a large peak at ∼118kDa, consistent with the dimer molecular weight, as opposed to the smaller molecular mass peak of the monomer (∼62kDa) (Fig. 7B and F). The FakA dimer was relatively stable, reaching an equilibrium with ∼57% of the protein in the dimer peak after 20 minutes (Fig. 7B and F).

**Figure 7:**
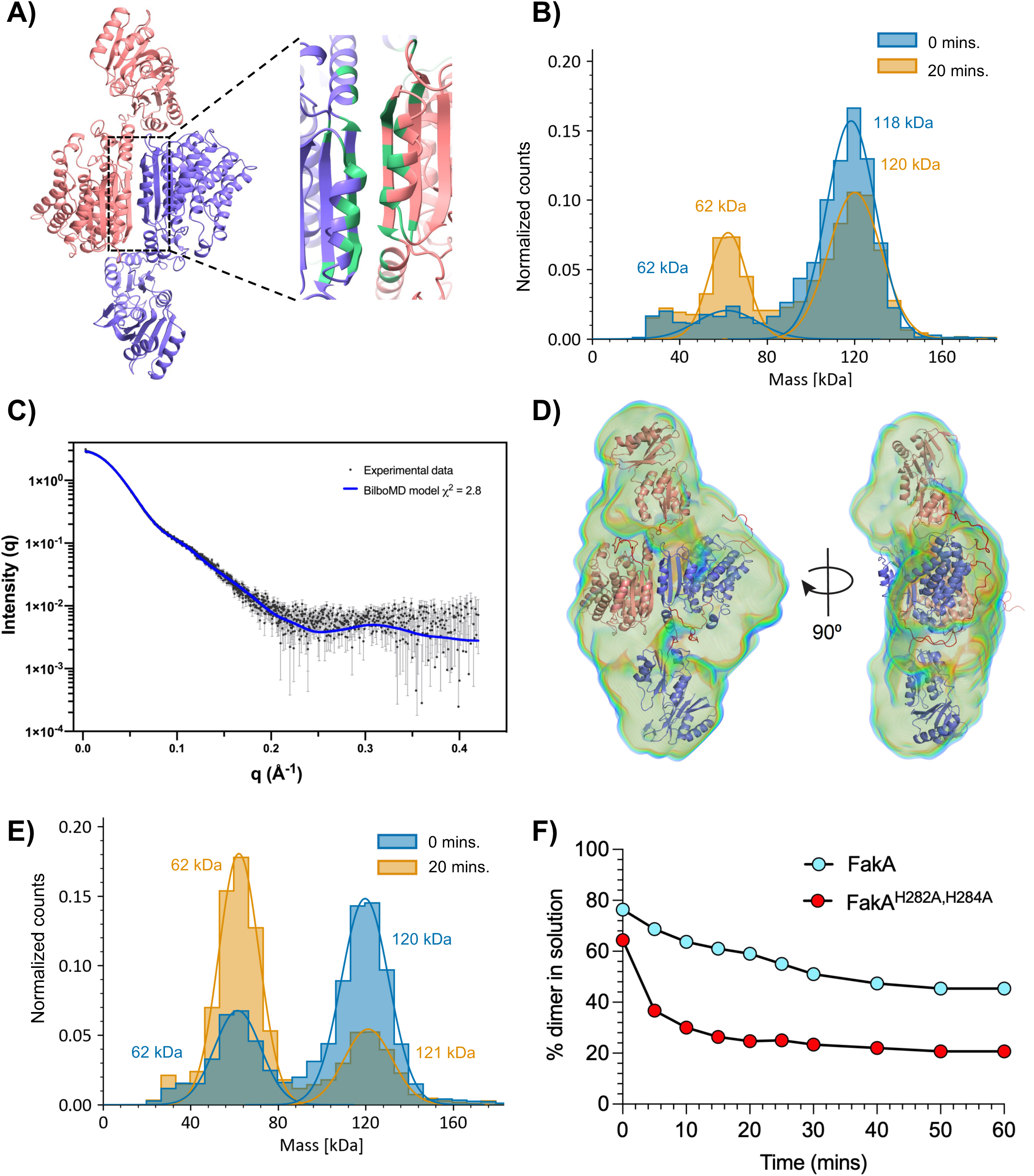
FakA is a homodimer mediated by the Middle domain in solution. **A)** AlphaFold model of the FakA homodimer. Predicted interacting residues are shown in green. **B)** Mass photometry of WT FakA. Measurements were taken at 0 (blue) and 20 minutes (orange). **C**) Crysol fitting of BilboMD models with Scattering intensity (I(q)) over the radial distance (q) experimental data. **D)** Molecular dynamics-based FakA model generated by BilboMD using the AlphaFold model (panel A) as an input overlayed within the electron density model of FakA using DENSS. The flexible regions are in red between rigid bodies (Purple and Pink). **E)** Mass photometry of FakA^H282A, H284A^ performed as described in panel B. **F)** Kinetic mass photometry showing the percent of wild-type in solution that is a dimer (light blue) compared to percent of FakA^H282A, H284A^ present as a dimer (red). Data represents the mean (n=3) with SEM.

To further investigate the dimerization and structure of FakA, we determined its size and shape by coupling size-exclusion chromatography with small-angle X-ray scattering (SEC-SAXS) (Fig. 7C-D, S8, Table S1 and S2). SAXS analysis revealed that FakA exists in solution as a dimer, with a radius of gyration (Rg) of 42.3 Å, and a maximum dimension of 160 Å (Table S1). The corrected Porod volume and Molecular weight estimated from SAXS data is also consistent with FakA forming a dimer (Table S1). Initial fitting of our predicted dimer AlphaFold model to the SAXS data did not give a good fit; however, it was greatly improved using BilboMD by flexible modeling of disordered residues at the linker regions connecting the N-terminal, Middle, and C-terminal domains (χ^2^ = 2.8; Fig. 7C-D, S8, Table S1, and S2). SAXS-derived molecular envelopes (bead-based models by DAMMIF and electron density models by DENSS) also show that FakA is in a dimeric form that is mediated by the Middle domain (Fig. 7D and S8).

Because dimerization was predicted to occur within the Middle domain, we hypothesized that our FakA^H282A, H284A^ protein, which doesn’t bind Zn, would have altered dimerization capacity and dynamics. Thus, the same kinetic mass photometry experiment was performed on the mutant FakA. At the zero time point, 63% of the protein was within the dimer peak (Fig. 7E and F). Equilibrium was again reached after approximately 20 minutes with only around 24% in the dimer form. Together, these data support a model whereby FakA forms a stable dimer in solution which is mediated by the Middle domain. Furthermore, it demonstrates that disruption of the metal binding pocket negatively impacts dimer formation and FakA function.

### FakA-FakB interaction

FakA has been shown to interact with the FakBs as part of the FAK complex ^26, 30^. Gel-filtration data of combined FakA and FakB2 proteins demonstrated that FakA and FakB2 are found in the same fraction as observed by SDS-PAGE (Fig. 8A). This is consistent with what has been observed in *Streptococcus* ^29^. To measure the affinity of the interaction of FakA with FakB2, we used bio-layer interferometry (Fig. 8B, and Table S3). His-FakB2 was immobilized on Ni-NTA biosensor tips and then exposed to varying concentrations of FakA. The results show a strong interaction with a dissociation constant (KD) of ∼4.34 nM when measured as a 1:1 ratio of FakA:FakB2 (R^2^ = 0.97 and a χ^2^ = 14.68, Table S3). Since the ratio and quantity of FakA and FakB in the FAK complex have not yet been elucidated, we envision two scenarios: one where the FakA dimer separates allowing for a FAK complex of a FakA/FakB heterodimer and a second being a heterotetrameric complex where the FakA dimer interface is maintained (Fig. 9A). AlphaFold-Multimer was used to predict multiple potential orientation outcomes: a heterodimer or a heterotetramer that could be formed with 2 FakA proteins and 2 FakB proteins (Fig. 9B-C, and S9). While the biological significance of each model is yet to be validated, each model orientates the headgroup of the fatty acid in FakB within the proximity of the ATP in the kinase domain of FakA. To account for the scenario that dimerized FakA potentially bound immobilized FakB2, the BLI data was re-calculated using bivalent analyte computation, giving a new KD of 6.65nM (Fig. 8B, and Table S3). This model fits the data more closely with an R^2^ value of 0.98 and a χ^2^ value of 2.55. This supports the hypothesis that FakA binds FakB2 with high affinity and that FakA is a dimer upon FakB2 binding.

**Figure 8:**
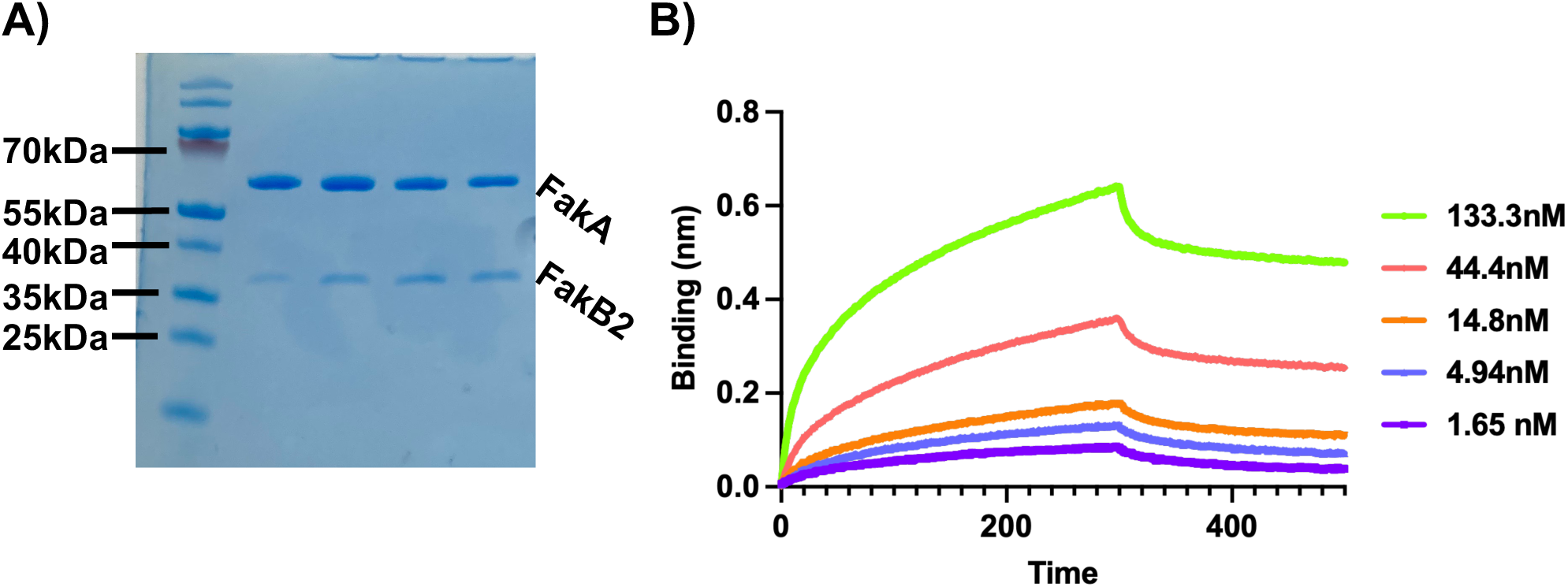
FakA and FakB2 Interact. **A)** SDS-PAGE of gel filtration showing co-elution of FakA and FakB2. **B)** BLI using sequential treatment of His-FakB2, 0.1 mg mL^-1^ BSA, buffer (wash), buffer (baseline), indicated concentration of FakA, and buffer (dissociation). Estimated KD 4.34 nM (1:1 analysis) or 6.65 nM (bivalent analyte analysis (1:2).

**Figure 9.**
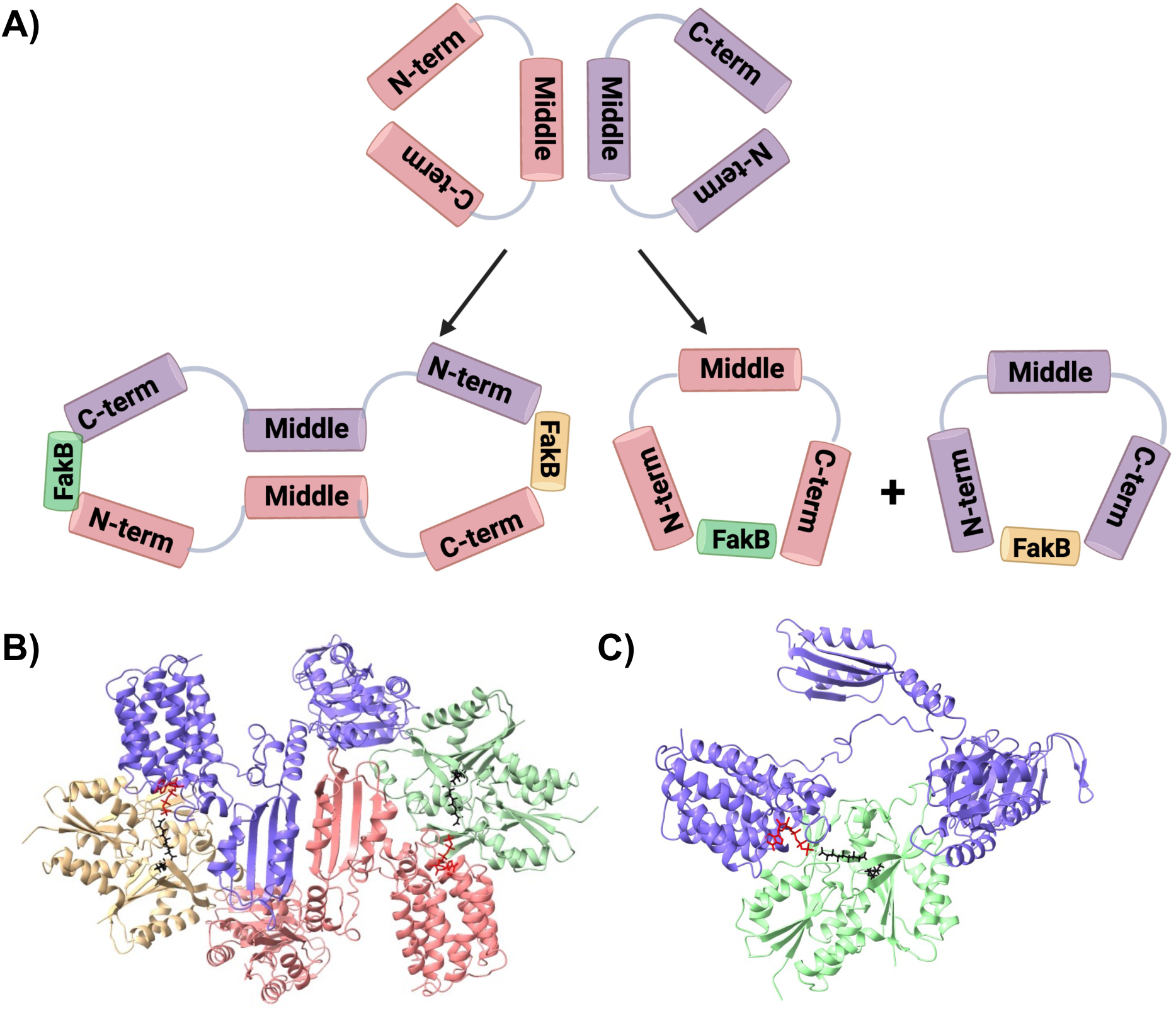
Models of FakA FakB2 interactions. **A)** Cartoon model of FakA-FakB potential interactions. FakA likely homodimerizes and either forms a heterotetramer or heterodimers with FakB. **B)** AlphaFold model of FakA-FakB2 heterotetramer. **C)** AlphaFold model of FakA-FakB2 heterodimer. AMP-PNP is rendered as red cylinders and oleic acid in black. Panel A made with bioRENDER.

## Discussion

While the FAK system’s importance in the use of exoFAs for lipid synthesis is known, little is known about the mechanism by which fatty acids are phosphorylated by the system or the identity of the composition of the complex. Our data, along with others, demonstrate that the FakA protein contains three specific domains: a catalytic N-terminal domain that phosphorylates a fatty acid, a Middle domain involved in dimerization, and a C-terminal domain with structural similarity to acyl-binding proteins like FakBs. We have provided context to kinase function during catalysis through a comparison of our newly solved Apo-FakA_N and AMP-PNP-FakA_N structures (Fig. 3). In particular, the α1-2 and α5-6 loops move to bind and secure ATP in preparation for the fatty acid-bound FakB binding in a way to position the γ-phosphate in proximity to the fatty acid head group. We have termed the α1-2 loop an activating loop since it 1) contains critical Asp residues that must reposition upon ATP binding for kinase activity 2) stabilizes that ATP molecule, and 3) would sterically hinder access by other proteins to the active site if it did not collapse around the ATP. We predict that this region of FakA could provide a regulatory mechanism to allow FakA-FakB binding when ATP and fatty acids are plentiful. Indeed, the collapse of the activating loop changes the face of FakA, exposing an altered interface containing conserved hydrophobic residues that we predict facilitate complex formation (Fig. S2 and S6).

In our studies, we determined a metal binding pocket within the Middle domain of FakA that is necessary for kinase function and FakA’s downstream effects on virulence and growth (Fig. 5 and 6). We provide evidence that this pocket is involved in the maintenance of the structure necessary for FakA dimerization (Fig. 7). Our work identifies and supports specific amino acids within this region (C240, H282, H284) that are essential for kinase function. This metal binding is indeed important for function in the bacteria since we show that downstream *in vivo* effects on α-hemolysin and resistance to toxic UFAs are abrogated when this pocket is mutated to contain non-reactive alanines (Fig. 6). These residues are 100% conserved among FakA from Gram-positive bacteria that we have investigated (Fig. S6). It is unlikely that these residues would comprise part of the active site. Instead, we predict that the Zn molecule stabilizes the β-sheets of the Middle domain, allowing proper folding. Based on the importance of zinc, it is tempting to speculate that there could also be a link between metal homeostasis and fatty acid metabolism. We show that FakA primarily homodimerizes in solution and our data supports the model that the Middle domain mediates this interaction (Fig. 7). We see two possible scenarios for the importance of this dimer. The first is that the FakA dimer is the “off state” and interaction with FakB occurs as a heterodimer following dissolution of the FakA dimer. Thus, FakA dimerization could be a regulatory mechanism. An alternate model is that FakA remains as a dimer during complex formation with FakB to form a heterotetramer. Dimerization of FakA may serve as an anchor for the complex and structurally support the binding of FakA and FakB. This would be particularly important in the transphosphorylation model predicted by AlphaFold (Fig. 9B). In this model, dimerization would be necessary for an active complex since the N-terminal domain of one FakA monomer interacts with FakB that is engaged with the C-terminal domain of the other FakA monomer. This could only happen if FakA remained as a dimer. Determining if FakA remains a dimer in the active FAK complex and whether the dimerization state serves a regulatory function will be one focus of future studies.

The studies by us and others provide the first insights into the structures of these ubiquitous proteins ^29, 30^; however, the details of how the phosphate is transferred from ATP to the fatty acid are yet to be elucidated. Our structures indicate that the activating loop movement may alter the face of FakA to facilitate access to the active site. We have shown that FakA and FakB directly interact with high affinity (Fig. 8). Solved structures of FakB1 and FakB2 bound demonstrate a channel in which the hydrophobic tail of the fatty acid is positioned ^25^. This renders the headgroup exposed. We propose a model whereby an interface of the FakA kinase domain and FakB form an active site to allow phosphotransfer of the ATP γ-phosphate in FakA is transferred to the fatty acid headgroup exposed by FakB. This model is unaffected by whether the complex exists as a heterodimer or heterotetramer (Fig. 9). This is similar to a recently proposed model for *S. aureus* FakA and FakB ^30^. Also consistent between our models and theirs is that the FakA C-terminal domain serves as a support for FakB, binding to the opposite side of FakB compared to the position of the fatty acid. This specific interaction provides additional stabilization to the interaction between FakA and FakB. Indeed, the C-terminal amino acid of FakA is important for interacting with FakB which we hypothesize stabilizes the complex ^30^. However, a second model was recently proposed based on data using the *Streptococcus suis* Fak proteins ^29^. In this model, FakB binds the C-terminal domain of FakA, causing a conformational change to release the fatty acid into a pocket formed by FakA_C and FakA_N for phosphorylation. While we cannot rule out this possibility, our data favors the former model. Distinguishing between these two models and obtaining the structure of a functional complex will be the focus of our future studies.

*S. aureus* is a prevalent multi-drug-resistant pathogen. One of our long-term goals is to use fatty acid metabolism as a potential target for therapeutic development. Indeed, FASII inhibitors are one drug class that has been suggested to serve as novel therapeutics with one of the most well-known being triclosan. However, resistance to FASII inhibitors is known and Gram-positive bacteria could bypass them via exoFA acquisition through the FAK complex. Thus, finding potential targets within the FAK complex is necessary to develop therapeutic inhibitors to be used in combination with FASII inhibitors, effectively shutting down any mechanism of providing fatty acids for lipid synthesis. A better understanding of the FakA-FakB complex structure and functional dynamics is necessary to comprehend the mechanism of exoFA utilization in Gram-positive bacteria. Our data as well as others are making progress toward this goal, however, much is left to be elucidated. Further studies of the active complex as well as more knowledge of the regulatory mechanisms of this protein are necessary. Elucidation of this FAK complex will provide insight into a conserved mechanism among Gram-positives as well as inform downstream medicinal chemistry and molecular docking to create effective therapeutics and combat antibiotic resistance.

## Material and methods

### Bacterial strains, media, growth conditions, and cloning

The strains and plasmids used in this study are provided in Table S4. *S. aureus* strains were grown in tryptic soy broth or agar (TSB or TSA) supplemented with chloramphenicol (10 μg mL^-1^) or erythromycin (5 μg mL^-1^) when necessary. *Escherichia coli* strains were grown in lysogeny broth (LB) or 2X-YT medium with ampicillin (100 µg ml^-1^) or kanamycin (50 µg ml^-1^) when necessary. PCR reactions were performed with KOD Polymerase (Novagen, San Diego, CA), and products were cleaned using the ZymoResearch Clean and Concentrator kit (Irvine, CA) between steps. All inserts were sequenced at ACGT, Inc. (Wheeling, IL) to ensure no unintended changes occurred. Oligonucleotides are provided in Table S5.

### Construction of FakA variant expression plasmids for *S. aureus*

A PCR splicing by overlap extension (SOEing) technique was used to construct a *fakA^C^*^240^*^A^* variant. PCR amplification using oligos JBHEM1 and JBKU81, and JBKU82 and CNK26 were performed using pCK13 as a template. The products from those reactions were used in a SOEing PCR with JBHEM1 and CNK26. The final PCR product was digested with BamHI and PstI and cloned into the same sites of pCM28 to create pJB1036. A similar process was used for the *fakA^H^*^282^*^A, H^*^284^*^A^* variant. The first two PCR products used primer pairs JBHEM1/MP11 or CNK26/MP7 were mixed and used in a SOEing PCR, with subsequent cloning into pCM28 digested with BamHI and PstI to generate pMP4.

### Construction of expression plasmids for recombinant protein purification

The gene encoding FakA^D38A, D40A^ was amplified from pCK2 by PCR using primers CNK24 and CNK25 and cloned into pET28a digested with Nde1 and Xho1 such that a N-terminal His-tag was added to produce pCK10. For His-FakB2, the *fakB2* gene was amplified from AH1263 chromosome DNA using primers CNK39 and CNK40 and cloned into the NdeI and XhoI sites of pET28a to generate pCK23. The constructs containing the FakA Middle and C-terminal domains were amplified by PCR using primers JBKU104 and JB42 with pJB165 as the template and similarly cloned into pET28a to produce pJB1042. The constructs containing just the FakA C-terminal domain (pMJM2-4) as well as GST-TEV-tagged FakA WT (pMJM2), *fakA^H^*^282^*^A, H^*^284^*^A^* (pMJM3) and *fakA^C^*^240^*^A^* (pMJM4) in pET42A (+) were ordered from Genscript.

### Kinase Activity Assay

The Universal Kinase Kit (R and D systems. Cat #: EA004) was used following adjusted manufacturer instructions. The substrate mix was prepared in wells of a 96-well plate in a minimum of duplicate by combining 10 μL of ATP (1 mM) (or ADP for positive control) with 15 μL oleic acid (31.4 mM in MeOH) in each well. The enzyme mix was prepared by combining 10 μL of coupling phosphatase (5 ng μl^-1^), with 7.5 μL FakA (8 μM) and 7.5 μL FakB2 (16.14 μM). The enzyme and substrate mixes were combined in each well and the reaction was incubated at room temperature for 30 minutes with a control well of only Assay Buffer. 30 μL Malachite Green Reagent A, 100 μL diH2O, and 30 μL Malachite Green Reagent B were subsequently added, tapped to mix, then incubated at room temperature for 20 minutes. A620 was measured using a Tecan Spark 10M plate reader to determine phosphate production. This assay tracks inorganic phosphate levels. In short, the kinase activity of the Fak complex converts ATP to ADP which is subsequently converted to AMP and inorganic phosphate by a coupling phosphatase. It is the inorganic phosphate that interacts with malachite green for the colorimetric change measured at 620 nm. In all assays, an ADP control was used as a control for assay function.

### Inductively coupled plasma mass spectrometry (ICP-MS)

Purified proteins were diluted to a concentration of 1 mg mL^-1^ in Gel filtration buffer (50 mM Tris HCl [pH 7.4], 150 mM KCl, 1 mM TCEP, 5% glycerol in diH2O). Protein samples were compared to a control obtained as the flow-through of buffer through a 10 kDa concentrator during sample preparation and termed buffer control. Samples were provided to the K-State Veterinary Diagnostic Laboratory and a Trace Mineral Panel was performed using a Perkin Elmer NexIon 350 ICP-MS instrument.

### Hemolysis assay

Hemolysis assays were performed as previously described ^8^. Briefly, overnight cultures were diluted to an OD600=1, and 1 μL was spotted on TSA with 5% rabbit blood. Hemolysis activity was observed after 48 hours of incubation at 37°C.

### Bio-layer interferometry (BLI) for FakA-FakB2 interaction

BLI was performed on the Octet RED96e in the Kansas Intellectual and Developmental Disabilities Research Center at the University of Kansas Medical Center. Ni-NTA biosensor tips (Sartorius, Göttingen, Germany) were hydrated in Assay Buffer (50 mM Tris HCl [pH 7.4], 150 mM KCl, 1 mM TCEP, 5% glycerol, 10 mM MgCl2) for 10 minutes at room temperature. A baseline measurement was taken in buffer for 1 minute followed by loading/immobilization of 2 μg mL^-1^ purified His-tagged FakB2 protein for 4 minutes. The tips were then blocked with 0.1 mg mL^-1^ Bovine Serum Albumin (BSA) for 2 minutes followed by a 2-minute wash in Assay Buffer. A second baseline measurement was taken in Assay Buffer for 1 minute. Varying concentrations of untagged FakA purified protein were then allowed to bind to the immobilized FakB2 for 5 minutes followed by a 10-minute dissociation period in Assay Buffer. Data was used to determine an approximate KD.

### Mass Photometry (MP)

MP experiments were performed on a Refeyn TwoMP (Refeyn Ltd.) at the University of Iowa Protein & Crystallography Facility. Microscope coverslips (No.1.5H, 24 mm x 50 mm, Thorlabs Inc.) were cleaned by serial rinsing with Milli-Q water and HPLC-grade isopropanol (Sigma Aldrich), on which a CultureWell silicone gasket (Grace Bio-labs) was then placed. All MP measurements were performed at room temperature in buffer (50 mM Tris [pH 7.4], 150 mM KCl, 1 mM TCEP, 10 μM ZnSO4). For each measurement, 15 µL of buffer was placed in the well for focusing, after which 5 µL of 40 nM protein was introduced. Movies were recorded for 60 seconds at 50 fps under standard settings. MP measurements were calibrated using an in-house prepared protein standard mixture: β-Amylase (56, 112, and 224 kDa), and Thyroglobulin (670 kDa). MP data were processed using DiscoverMP (Refeyn Ltd).

## Protein Purification and Structural Analysis

Full details are provided in the Supplemental Materials.

## Data Availability

The coordinates and structure factors for Apo-FakA_N (8VIR), ADP-FakA_N (8VIT), Mn-FakA_N (8VIP), and AMP-PNP-FakA_N (8VIQ) have been deposited to the Worldwide Protein Databank (wwPDB). The SAXS data for full-length FakA have been deposited in Small Angle Scattering Bilogical Data Bank (SASBDB) under the accession code SASDTW8.

## Acknowledgments

This work was supported by NIH applications AI121073 and AI153773 to JLB and a Madison and Lila Self Graduate Fellowship to MJM. Additional support came from the University of Kansas COBRE-PSF NIH P30 GM110761. The use of the IMCA-CAT beamline 17-ID at the Advanced Photon Source was supported by the companies of the Industrial Macromolecular Crystallography Association through a contract with Hauptman-Woodward Medical Research Institute. Use of the Advanced Photon Source was supported by the U.S. Department of Energy, Office of Science, Office of Basic Energy Sciences under contract no. DE-AC02-06CH11357.

We would like to thank Jesse Hopkins and Maxwell Watkins at BioCAT beamline 18-ID-D (APS) for SEC-SAXS data collection. This research used resources of the Advanced Photon Source; a U.S. Department of Energy (DOE) Office of Science User Facility operated for the DOE

Office of Science by Argonne National Laboratory under Contract No. DE-AC02-06CH11357. BioCAT was supported by grant P30 GM138395 from the National Institute of General Medical Sciences of the National Institutes of Health. The content is solely the responsibility of the authors and does not necessarily reflect the official views of the National Institute of General Medical Sciences or the National Institutes of Health. We would also like to thank Rodrigo Jacamo at Sartorius for assistance with BLI experimental design and analysis.

## Author contributions

All authors made substantial contributions to the work through experimental design, data acquisition, data analysis and interpretation, or manuscript preparation and editing. All authors have approved the paper for submission and agree to be held accountable for the author’s own contributions. M.J.M., Z.X., and S.L. contributed to all aspects listed above; B.J.R. contributed to data acquisition, analysis, and interpretation; Z.R.D., M.J.R., C.N.K., T.S.F., M.M.K., and K.P.B. contributed to data acquisition; D.K.J., N.S., B.D.F., and J.L.B. contributed to the experimental design, data analysis and interpretation and manuscript preparation.

## Materials & Correspondence

Correspondence and material requests should be addressed to Jeffrey L. Bose.

## Statistical Information

Statistical analyses were performed using GraphPad Prism v.9 by Welch’s t-test or one-way ANOVA as described in the figure legends. Sample sizes are listed in figure legends. Error bars represent the Standard Error of the Mean (SEM).

---

**Figure S1: N-terminal FakA crystal structure hydrogen bonds. A)** Hydrogen bonds between nucleobase and sugar of the nucleotide and AMP-PNP-FakA_N. **B)** Hydrogen bonds between the triphosphate portion of the nucleotide and AMP-PNP-FakA_N. Hydrogen bonds are shown as blue dotted lines. Mg^2+^ ions are rendered as green spheres. AMP-PNP is rendered as gray cylinders and colored by element.

**Figure S2: N-terminal FakA electrostatic potential at FakB binding interface. A)** Electrostatic potential map of Apo-FakA_N. **B)** Electrostatic potential map of AMP-PNP-FakA_N. Mg^2+^ ions are rendered as green spheres. AMP-PNP is rendered as gray cylinders and colored by element. Surface shown on a scale from red (negative) to blue (positive).

**Figure S3: Comparison of ADP-FakA_N and AMP-PNP-FakA_N structures. A-C)** Close-up of ADP/AMP-PNP binding pocket. **D-F)** Full crystal structure of AMP-PNP-FakA_N, ADP-FakA_N, and an overlay. AMP-PNP-FakA_N is rendered in light blue, ADP-FakA-N in salmon, AMP-PNP and Mg^2+^ in red, and ADP and Mg^2+^ in black.

**Figure S4: C-terminal domain of FakA.** X-ray crystal structure of FakA_C

**Figure S5: Overlay of AlphaFold predicted model with solved crystal structures.** Apo-FakA_N in purple, FakA_C in yellow, AlphaFold model in gray.

**Figure S6: Conservation of key FakA amino acids.** CLUSTAL Omega 1.2.4 multiple sequence alignment of FakA from several Gram-positive bacteria through *S. aureus* FakA Pro288. Pink: residues are involved in Mg^2+^ coordination. Blue: α1-2 or α5-6 loop. Yellow: conserved hydrophobic region. Red: residues in Zn-binding pocket. Green: other noted residue in main text.

**Figure S7: Growth curve of FakA variants. A)** Growth of indicated strains in TSB variants without linoleic acid. **B)** Growth of indicated strains in TSB without or with 192 µM (0.006%) linoleic acid. Symbols represent that mean (n=3) with SEM. Error bars are present and may be smaller than symbols.

**Figure S8. Additional SAXS data and models.** SAXS-derived bead-based modeling of FakA by DAMMIF overlayed with BilboMD models. Four conditions (two symmetry and two calculation modes) were used. **A)** P1 fast, **B)** P1 slow, **C)** P2 fast, and **D)** P2 slow.

**Figure S9. Alternative Models of FakA FakB interactions. A)** Alternative AlphaFold model of FakA FakB2 heterotetramer. **B)** AlphaFold model of FakA FakB1 heterodimer. AMP-PNP is rendered as red cylinders and oleic acid in black.

**Movie S1.** Chimera morph video of Apo-FakA_N to AMP-PNP-FakA_N in full view. Mg^2+^ ions are rendered as green spheres. AMP-PNP is rendered in gray cylinders with important atoms colored by element.

**Movie S2.** Chimera morph video of Apo-FakA_N to AMP-PNP-FakA_N showing nucleobase and sugar binding pocket of FakA_N. Mg^2+^ ions are rendered as green spheres. AMP-PNP is rendered in gray cylinders with important atoms colored by element.

**Movie S3.** Chimera morph video of Apo-FakA_N to AMP-PNP-FakA_N showing triphosphate binding of FakA_N. Mg^2+^ ions are rendered as green spheres. AMP-PNP is rendered in gray cylinders with important atoms colored by element.

**Movie S4.** Chimera morph video of Apo-FakA_N to AMP-PNP-FakA_N showing Mg^2+^ coordination by FakA_N. Mg^2+^ ions are rendered as green spheres. AMP-PNP is rendered in gray cylinders with important atoms colored by element.

**Movie S5.** Chimera morph video of Apo-FakA_N to AMP-PNP-FakA_N showing hydrophobic face of FakA_N for FakB interaction. Mg^2+^ ions are rendered as green spheres. AMP-PNP is rendered in gray cylinders with important atoms colored by element. Hydrophobic residues are rendered in dark green.

