## Supplementary Materials and Methods, Fig. S1 to S9, Table S1 to S7 for "Molecular basis for the activation of the Fatty Acid Kinase complex of *Staphylococcus aureus*"

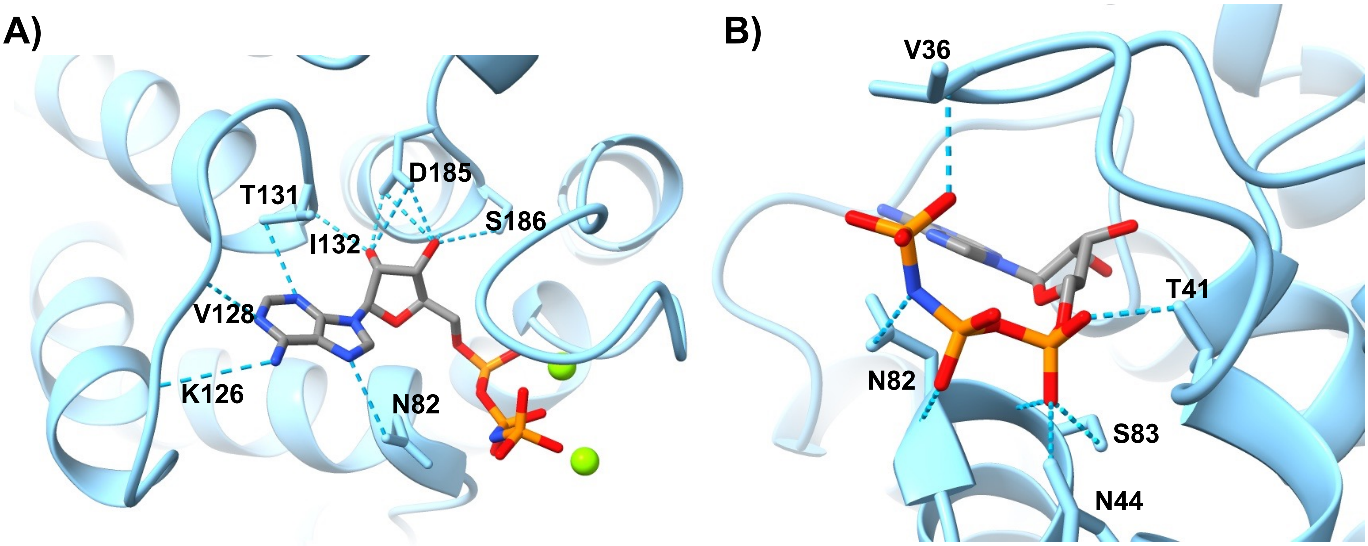


**Figure S1: N-terminal FakA crystal structure hydrogen bonds. A)** Hydrogen bonds between nucleobase and sugar of the nucleotide and AMP-PNP-FakA_N. **B)** Hydrogen bonds between the triphosphate portion of the nucleotide and AMP-PNP-FakA_N. Hydrogen bonds are shown as blue dotted lines. $Mg^{2+}$ ions are rendered as green spheres. AMP-PNP is rendered as gray cylinders and colored by element.

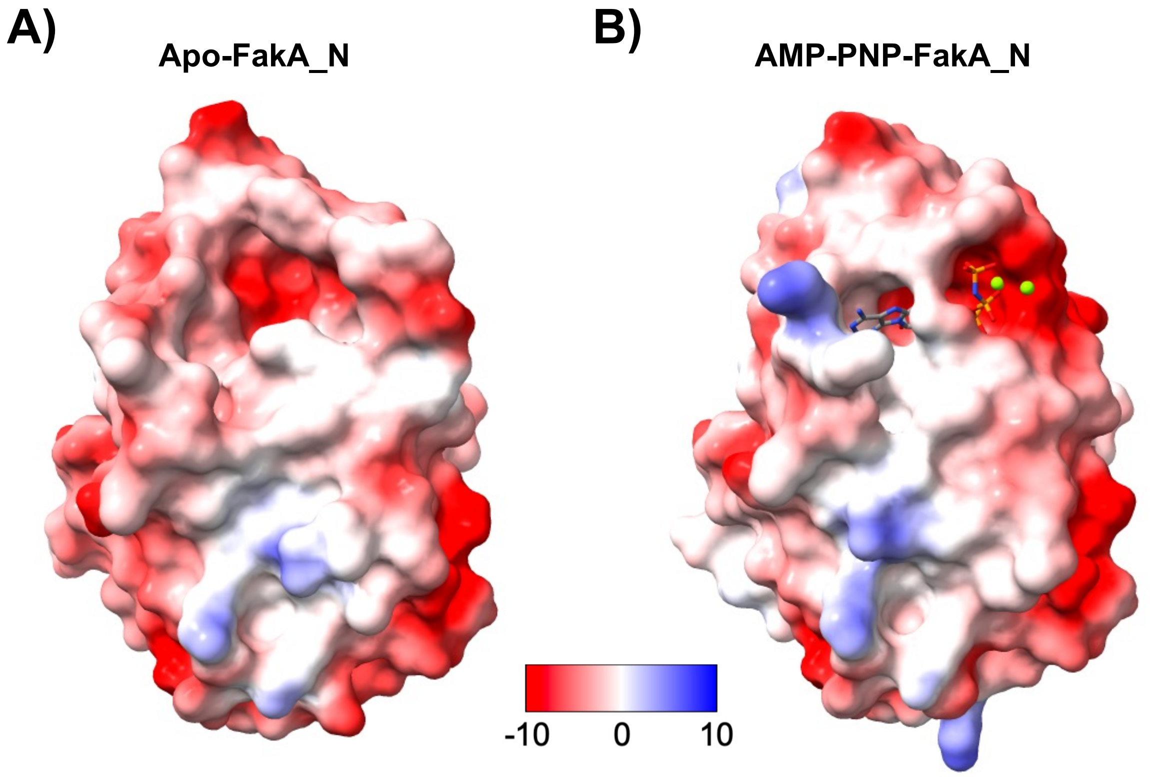


**Figure S2: N-terminal FakA electrostatic potential at FakB binding interface. A)** Electrostatic potential map of Apo-FakA_N. **B)** Electrostatic potential map of AMP-PNP-FakA_N. $Mg^{2+}$ ions are rendered as green spheres. AMP-PNP is rendered as gray cylinders and colored by element. Surface shown on a scale from red (negative) to blue (positive).

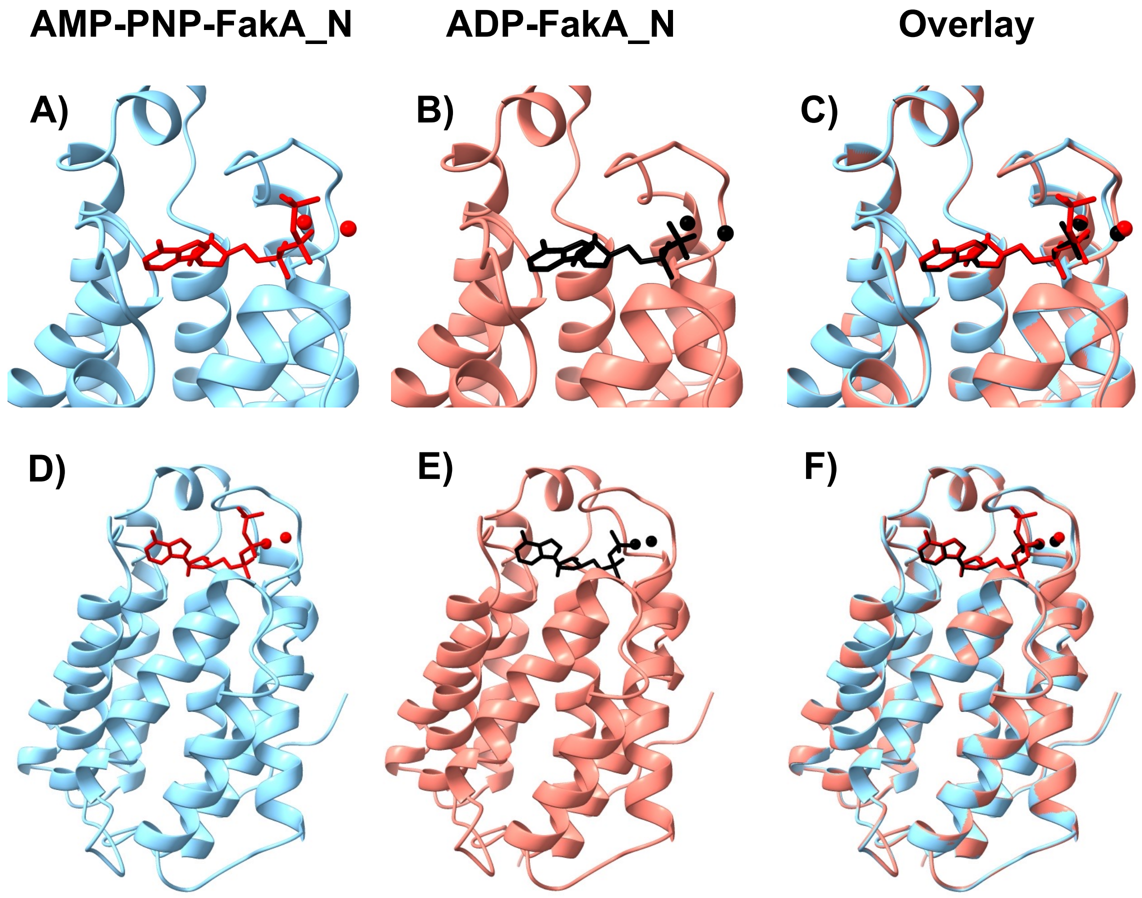


**Figure S3: Comparison of ADP-FakA_N and AMP-PNP-FakA_N structures. A-C)** Close-up of ADP/AMP-PNP binding pocket. **D-F)** Full crystal structure of AMP-PNP-FakA_N, ADP-FakA_N, and an overlay. AMP-PNP-FakA_N is rendered in light blue, ADP-FakA-N in salmon, AMP-PNP and $Mg^{2+}$ in red, and ADP and $Mg^{2+}$ in black.

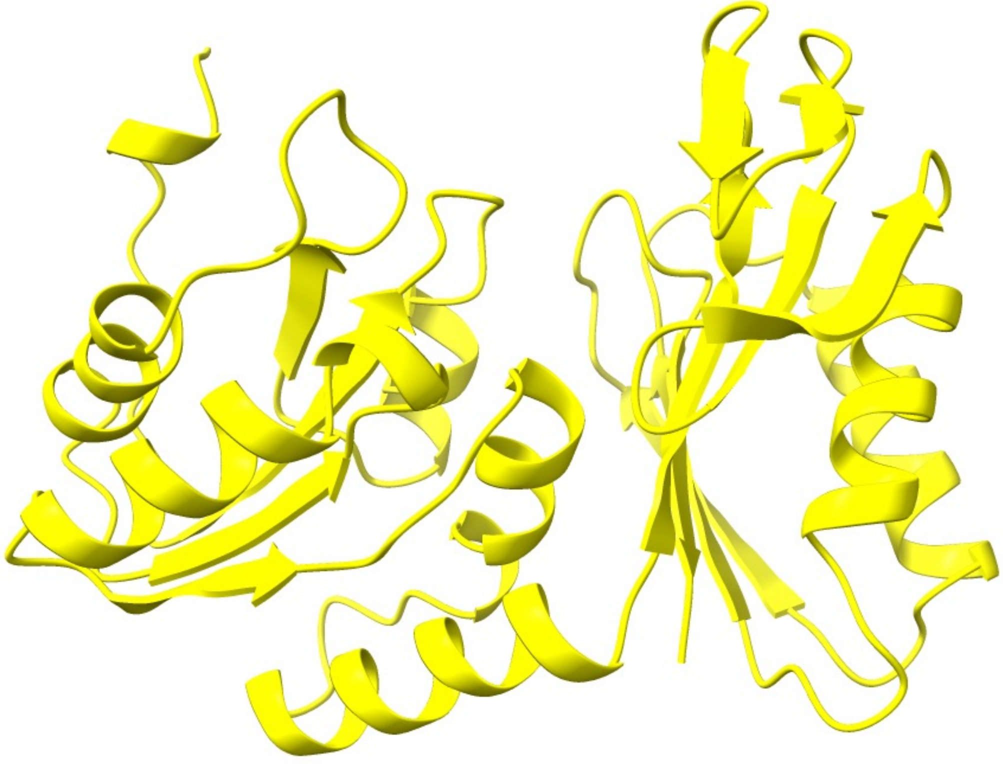

**Figure S4: C-terminal domain of FakA.** X-ray crystal structure of FakA_C

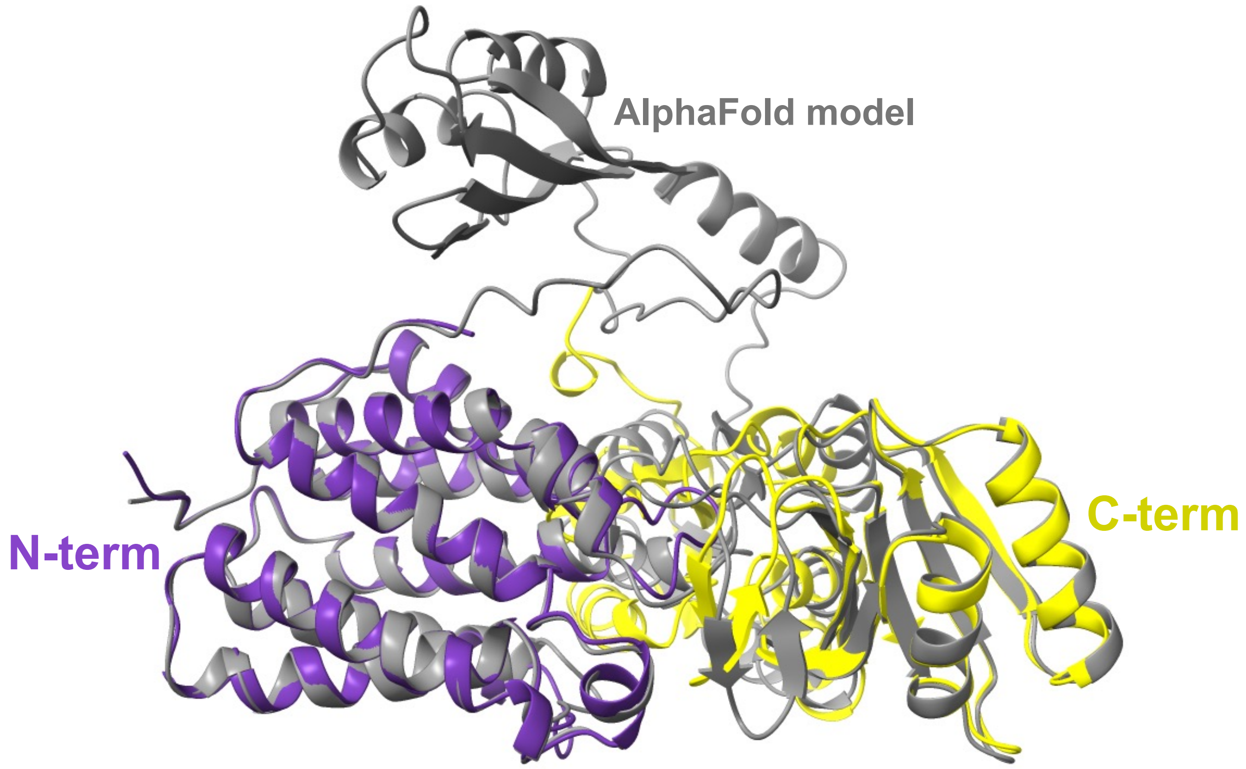


**Figure S5: Overlay of AlphaFold predicted model with solved crystal structures.** Apo-FakA_N in purple, FakA_C in yellow, AlphaFold model in gray.

```
                    N-terminal Domain
S.aureus FakA       ---MISKINGKLFADMIIQGAQNLSNNADLVDSLNVYPVPDGDTGTNMNLTMTSGREEVE
CP1732              --MKFNKVNGEHLYYMMSNAANKLELQSEYVNSLNVFPVPDGDTGTNMSMTFRAAVKEIE
Listeria            --MSIYQLDPEKFAAMIALGAENLAKNADFVDSLNVFPVPDGDTGTNMNLSMTSGAEEVA
PEPE_0838           MEAIGTLITNEVFDQMVQASSDRLKMNADFINSLNVFPVPDGDTGTNMSLSLESGAKYEN
LBA1311             --MVINEIDSKQFRDMVRVATHRMGKNAEFVNSLNVFPVPDGDTGTNMNLTIESGAKAVS
EF_3114             --MNVTEISAGQFQEMVQAGATRLHTNAEYVNSLNVFPVPDGDTGTNMNLSMTSGAKAVA
L193593             -----------------------MNEQAEYVNSLNVFPVPDGDTGTNMGMTITNGAKDVA
SP_0443             ----MSKITTSLFQEMVQAASTRLNKQAEYVNSLNVFPVPDGDTGTNMGMTIENGAKEVA
SMU_229             ----MANITTSLFQGMVQAAATRLGKQAEYVNSLNVFPVPDGDTGTNMGMTMDNGAKEVS
SPy_1886            ----MSNITTSLFQEMVQAAATRLGKQAEYVNSLNVFPVPDGDTGTNMSMTMDNGAKEVA
BSU15840            --MSIRTLDGRTFAEMILAGAQNLSQNASAVDALNVFPVPDGDTGTNMNLSMTSGAREVE
                                           :  ::. :::***:***********.:::  . .

S.aureus FakA       NNLSKNIGELGKTFSKGLLMGARGNSGVILSQLFRGFCKNIESESEINSKLLAESFQAGV
CP1732              GMDNKNIGEVSKKLAKGALMGARGNSGVILSQILRGISKGLEGKEEADASEFANALLEGS
Listeria            KNDKETISAVGANLAKGLLMGARGNSGVILSQLFRGFSKSIENKETLNAEEFANAFVKGV
PEPE_0838           DDASLEVGHLAASLAKGLLMGARGNSGVILSQIFRGFAKSVEGKATLNAEDLAEAVAEGA
LBA1311             ENPSTSVGDLTESLAKGMLMGARGNSGVITSQLFRGFYKATQGMKTLTAPELANAFSNGV
EF_3114             DSRSEKVGELAGALSKGLLMGARGNSGVILSQLFRGFSKQIPEVTTLNAKDLAAAFTHGV
L193593             SKPADTVGQVAQILSKGLLMGARGNSGVILSQLFRGFAQYVKDYEEIDGIHLAAALQNGV
SP_0443             DKPASTVGEVASILAKGLLMGARGNSGVITSQLFRGFSQAIKDKDELTGQDLALAFQSGV
SMU_229             DKPASTVGEVGQILSKGLLMGARGNSGVITSQLFRGFGQSIKNKEELTGKDLAQALQAGV
SPy_1886            DKPASTVGEVGQMLSKGLLMGARGNSGVITSQLFRGFGQSIKGKDELTGKDLAQAFQVGV
BSU15840            QMDTDDIGKVGSALSKGLLMGARGNSGVILSQLFRGFSKNIETKKEINALEFAAALQAGV
                          :. :   ::** *********** **::**: :         .  :* :.  *

S.aureus FakA       ETAYKAVMKPVEGTILTVAKDAAQAAIEKANNTEDCIELMEYIIVKANESLENTPNLLAV
CP1732              KSAYKAVMRPTEGTILTIVRTAGEVAV--ALKEDNITELMREVCKESKIMLDKTPEMLPA
Listeria            ETAYKAVMKPVEGTILTVAREAAKAGVLAANEHQEIELVMEAILKQGKVALEKTPDLLPV
PEPE_0838           KTAYKAVMKPTEGTILTVVREAAKAGNEAIKTTDSLAEVMEAIDSAAQTALKKTTDLLPV
LBA1311             ATAYKAVMKPVEGTILTVARVAAQEGTSKANDTDDVTEVMKAVVEGAKKALKSTPDLLPV
EF_3114             ETAYKAVMKPVEGTILTVSREAARSGERKAKETDDCIEVMTAVVAGAKRALAKTPDLLPV
L193593             EVAYKAVMKPVEGTILTVSRGAAELAKRKTDETDDAVKIMEAALEGAKKALAMTPDMLPV
SP_0443             EVAYKAVMKPVEGTILTVSRGAAIGAKKKAEQTDDAVEVMRAALEGAKTALAKTPDMLPV
SMU_229             EVAYKAVMKPVEGTILTVSRGAATAAIKKSENTDDTVEVMQAALDGAKAALAKTPDLLPV
SPy_1886            EVAYKAVMKPVEGTILTVSRGAATAALKKADLTDDAVEVMQAALDGAKGALAKTPDLLPV
BSU15840            DMAYKAVMKPVEGTILTVAKDAAKKAMILAEKETDITALMTAVTEEAEASLNRTPELLPV
                      ******:*.******: : *.  .        .   :*      .:  *  * ::* .
                                                 Unstructured region
S.aureus FakA       LKEVGVVDSGGKGLLCVYEGFLKALKGEKVEAKVAK---IDKDEFVHDEH---DFHGVIN
CP1732              LKKAKVVDSGGMGLLIILQGMQEALENGLKVTTGT---PQAVK----SSVAKAQRSETMS
Listeria            LKEVGVVDSGGQGLIIIYEGFYGVLTGKMAEAPDYM---ASMGELINAEHHR-HVQDFMS
PEPE_0838           LKEVGVVDSGGQGLVFVFDAFNDVLNGNDITEYKHEPDATEVNEMVDAKHHQ-SVQGKIN
LBA1311             LKQVGVVDSGGQGLVFIYEGFLEGLLGENFA-DQYQPDENEMDEMINAMHHQSSVQSQLA
EF_3114             LKEVGVVDSGGQGLLFIYEGFLEALSGEFLPTETYEPSPAEMDEMVNAEHHR-GVSGHVA
L193593             LKEVGVVDSGGQGLVYIYEGFLMALNGEFVPETPV-AELGAMDRMVNVEHES---VANAS
SP_0443             LKEVGVVDSGGQGLVFIYEGFLSALTGEYIASEDFVATPANMSEMINVEHHK-SVAGHVA
SMU_229             LREVGVVDSGGQGLVFIYEGFLSALTGDYIISEDFQATPANMTEMINAEHHK-SVASHVA
SPy_1886            LKEVGVVDSGGQGLVFIYEGFLSALNGDYVTSADFKATPANMSEMINAEHHK-SVVGHVA
BSU15840            LKEVGVVDSGGKGLLCVYEGFLASLKGETVPQKAVL---PSLDDMVSAEHHK-SAQSMMN
                    *::. ****** **: : :.:   * .
                      Middle domain
S.aureus FakA       TEDIIYGYCTEMMVRFGKN---KKAFDEQEFRQDMSQFGDSLLVINDEEIVKVHVHTEYP
CP1732              EEDIKFGYCTEFIILGDS-------NHAEEFKSKVINKGDSLVVVGYEDVIKVHIHTNNP
Listeria            TEDIHFGYCTEIIVKINENKPGLKPFDEEQFRQDLSELGDSLLVAADDEVVKVHVHVEHP
PEPE_0838           PEDIEFGYCTQMMVRIGKGLQVTKEFDYDTFYNYLAKMGDSLLVINDDEVVKVHVHTEHP
LBA1311             TQDIKNGYCTEIMVDLNADVPNKKEFDLEEFRKHLSGLGDSLLAVSDGEIAKVHVHTEHP
EF_3114             TEDIKFGYCTEIMVRIGEGPTVDSAFDYETFRNYLNELGDSLLVVNDDEIIKVHVHTEHP
L193593             TADIKFGYCTEIMVELGKGPTSRESYDHDNFQAYLAGIGNSLLVVDDEEVVKVHVHTEDP
SP_0443             TEDITFGYCTEIMVALKQGPTYAKDFDYDEFRNYLDELGDSLLVVNDDEIVKVHVHTEDP
SMU_229             TEDITFGYCTEIMVALKQGPTYVKDFNYENFRNYLNNLGDSLLVVNDDEIVKVHVHTEDP
SPy_1886            TEDITYGYCTEIMVALKQGPTYVKEFNYDEFQGYLSGLGDSLLVVNDDEIVKVHVHTEDP
BSU15840            TEDIEFGFCTEVMVRLDQT---KREFDEGTFRQDLSQFGDSLLVIADESLAKVHIHAEEP
                      **  *:**:.::            .   *   :   *:**:.    .: ***:*.: *
```

**Figure S6: Conservation of key FakA amino acids.** CLUSTAL Omega 1.2.4 multiple sequence alignment of FakA from several Gram-positive bacteria through *S. aureus* FakA Pro288. Pink: residues are involved in $Mg^{2+}$ coordination. Blue: α1-2 or α5-6 loop. Yellow: conserved hydrophobic region. Red: residues in Zn-binding pocket. Green: other noted residue in main text.

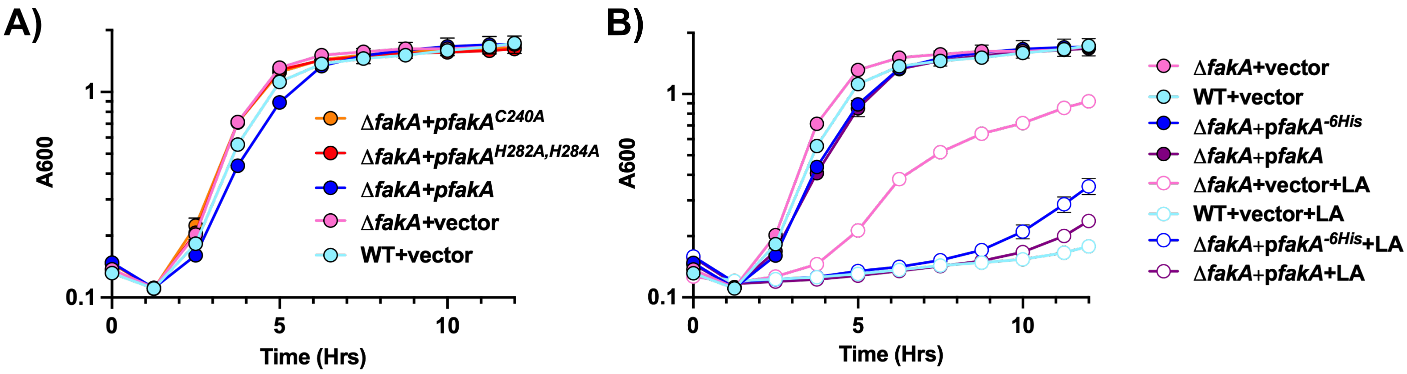


**Figure S7: Growth curve of FakA variants. A)** Growth of indicated strains in TSB variants without linoleic acid. **B)** Growth of indicated strains in TSB without or with 192 μM (0.006%) linoleic acid. Symbols represent that mean (n=3) with SEM. Error bars are present and may be smaller than symbols.

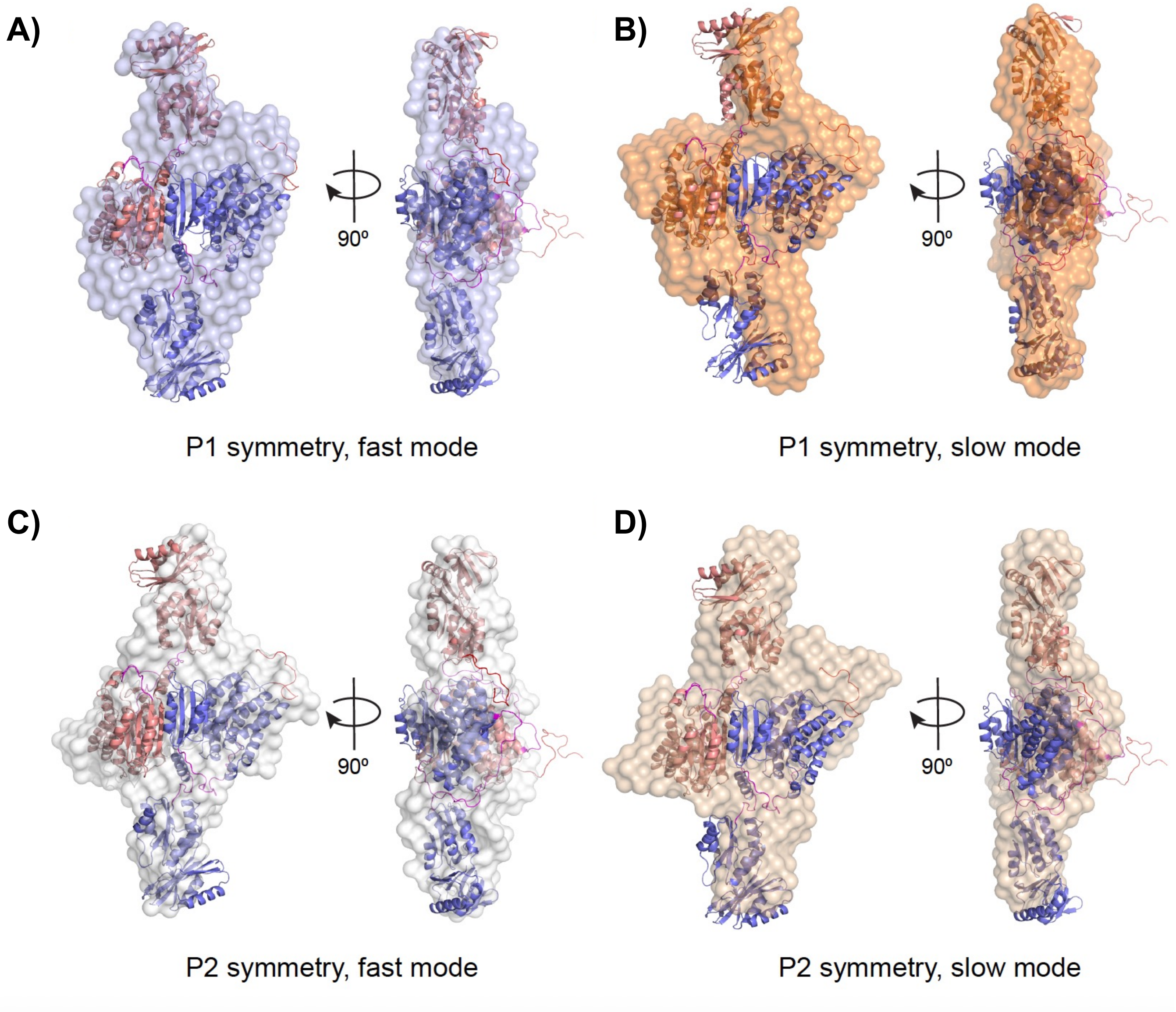


**Figure S8. Additional SAXS data and models.** SAXS-derived bead-based modeling of FakA by DAMMIF overlayed with BilboMD models. Four conditions (two symmetry and two calculation modes) were used. **A)** P1 fast, **B)** P1 slow, **C)** P2 fast, and **D)** P2 slow.

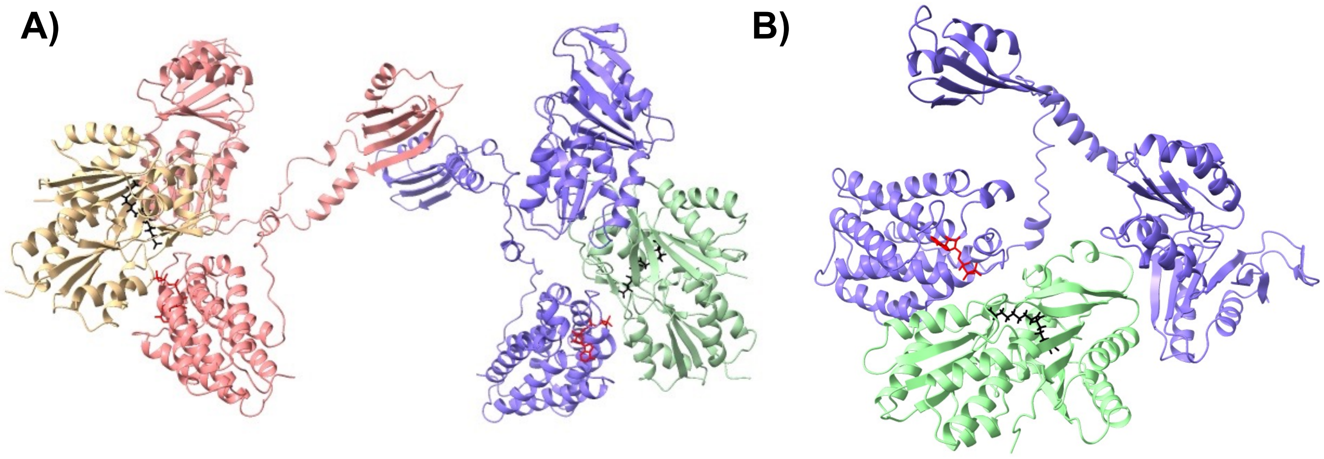


**Figure S9. Alternative Models of FakA FakB interactions. A)** Alternative AlphaFold model of FakA FakB2 heterotetramer. **B)** AlphaFold model of FakA FakB1 heterodimer. AMP-PNP is rendered as red cylinders and oleic acid in black.

**Movies S1-5.**

**Movie S1.** Chimera morph video of Apo-FakA_N to AMP-PNP-FakA_N in full view. $Mg^{2+}$ ions are rendered as green spheres. AMP-PNP is rendered in gray cylinders with important atoms colored by element.

**Movie S2.** Chimera morph video of Apo-FakA_N to AMP-PNP-FakA_N showing nucleobase and sugar binding pocket of FakA_N. $Mg^{2+}$ ions are rendered as green spheres. AMP-PNP is rendered in gray cylinders with important atoms colored by element.

**Movie S3.** Chimera morph video of Apo-FakA_N to AMP-PNP-FakA_N showing triphosphate binding of FakA_N. $Mg^{2+}$ ions are rendered as green spheres. AMP-PNP is rendered in gray cylinders with important atoms colored by element.

**Movie S4.** Chimera morph video of Apo-FakA_N to AMP-PNP-FakA_N showing $Mg^{2+}$ coordination by FakA_N. $Mg^{2+}$ ions are rendered as green spheres. AMP-PNP is rendered in gray cylinders with important atoms colored by element.

**Movie S5.** Chimera morph video of Apo-FakA_N to AMP-PNP-FakA_N showing hydrophobic face of FakA_N for FakB interaction. $Mg^{2+}$ ions are rendered as green spheres. AMP-PNP is rendered in gray cylinders with important atoms colored by element. Hydrophobic residues are rendered in dark green.

**Table S1.** SAXS data analysis statistics

**a) Structural parameters**

| | |
|---|---|
| Guinier analysis | |
| $R_g$ (Å) | 42.70 ± 0.04 |
| *q*-range ($Å^{-1}$) | 0.00292-0.03039 |
| *q*max$R_g$ | 1.2975 |
| I(0) (Arb.) | 2.904 ± 0.002 |
| *Coefficient of correlation, $r^2$* | 0.9967 |
| *GNOM analysis* | |
| $R_g$ (Å)** | 43.34 ± 0.04 |
| Dmax (Å)** | 160 |
| I(0) (Arb.) | 2.913 ± 0.002 |
| *q*-range ($Å^{-1}$) | 0.00292-0.18742 |
| $\chi^2$ | 1.6731 |
| *Total estimate from GNOM)* | 0.851 |
| Volume ($Å^3$, adjusted $V_P$ as SAXS MoW2) | 179000 |
| MW, Vp method (kDa) | 148.4 |
| MW, Vc method (kDa) | 129.4 |
| MW, (S&S) (kDa) | 119.9 |
| MW, (Bayes) (kDa) | 118.8 |
| MW, Theoretical by dimer sequence (kDa) | 125.4 |

**b) bead-based modeling parameters**

| | | | | |
|---|---|---|---|---|
| DAMMIF (default parameters, 20 calculations) | | | | |
| q-range for fitting ($Å^{-1}$) | 0.00292-0.18742 | | | |
| Calculation mode | Fast | Slow | Fast | Slow |
| Symmetry, anisotropy assumptions | P1,unknown | P1,unknown | P2,unknown | P2,unknown |
| Ambiguity score (AMBIMETER) | 2.477 | | | |
| NSD (standard deviation) | 0.89 ± 0.11 | 1.10 ± 0.15 | 0.68 ± 0.11 | 0.75 ± 0.14 |
| $\chi^2$ range | 1.671-1.814 | 1.667-1.717 | 1.681-1.965 | 1.707-1.794 |
| Model MW estimate range | 141.55-145.54 | 143.93-145.62 | 143.29-147.29 | 143.71-146.76 |
| Model $R_g$ range | 43.3-43.3 | 43.3-43.3 | 43.3-43.4 | 43.3-43.3 |
| Model $D_{max}$ range | 162.4-180.5 | 162.6-176.5 | 161.8-181.9 | 166.9-177 |
| Resolution (SASRES) (Å) | 44.1 | 38.4 | 32.0 | 27.8 |
| DAMMIN (Refinement of damstart.pdb) | | | | |
| $\chi^2$ | 1.655 | 1.681 | 1.822 | 1.667 |
| Model MW estimate (ratio to expected dimer) | 153.34 (1.22) | 154.99 (1.24) | 143.38 (1.14) | 152.68 (1.22) |
| Model $R_g$ | 43.1 | 43.3 | 43.4 | 43.4 |

**c) DENSS modeling parameters** (default parameters, 20 calculations)

| | |
|---|---|
| q-range for fitting ($Å^{-1}$) | 0.00292-0.18742 |
| $\chi^2$ range | 0.0481-0.62658 |
| $R_g$ range (Å) | 41.33-42.33 |
| Support volume range ($Å^3$) | 421031-497172 |
| Real space correlation (RCS) | 0.7839 ± 0.0467 |
| $\chi^2$ (refine) | 0.03517 |
| $R_g$ (refine, Å) | 42.45 |
| Support volume (refine, $Å^3$) | 449297 |
| Reconstruction Resolution (Å) | 49.6 ± 11.0 |

**d) BILBOMD modeling parameters**

| | |
|---|---|
| Extent of conformational sampling | 400 conformations per Rg |
| $R_g$ range (Å) | 37-47 |
| $\chi^2$ | 2.847 |

**Table S2.** SAXS data collection statistics

**a) Sample details**

| | |
|---|---|
| SEC Column | Superdex 200 increase 10/300 GL |
| Loaded concentration (mg/ml) | 4.7 |
| Injection volume (ul) | 200 |
| Flow rate (ml/min) | 0.6 |
| Solvent (solvent blanks taken from SEC flowthrough prior to elution of protein) | 50 mM Tris pH 7.4, 150 mM KCl, 1 mM TCEP, 5% glycerol |

**b) SAXS data-collection parameters**

| | |
|---|---|
| Instrument | BioCAT facility at the Advanced Photon Source beamline 18ID with Pilatus3 X 1M detector |
| Wavelength (Å) | 1.033 |
| Beam size ($um^2$) | 150 (h) x 25 (v) focused on the detector |
| Camera length (m) | 3.663 |
| $q$ measurement range ($Å^{-1}$) | 0.0029-0.42 |
| Absolute scaling method | Glassy Carbon, NIST SRM 3600 |
| Basis for normalization to constant counts | To transmitted intensity by beam-stop counter |
| Monitoring for radiation damage | Automated frame-by-frame comparison of relevant regions using CORMAP (Franke et al., 2015) implemented in BioXTAS RAW |
| Exposure time | 0.5 s exposure time with a 1 s total exposure period (0.5 s on, 0.5 s off) of entire SEC elution |
| Sample configuration | SEC-SAXS with sheath-flow cell (Kirby et al., 2016), effective path length 0.542 mm. |
| Sample temperature (ºC) | 23 |

**c) Software employed for SAXS data reduction, analysis, and interpretation**

| | |
|---|---|
| SAXS data reduction | Radial averaging; frame comparison, averaging, and subtraction done using BioXTAS RAW 2.1.4 (Hopkins et al., 2017) |
| Basic analysis: Guinier, MW, Normalized Kratky, P(r) | Guinier fit and M.W. using BioXTAS RAW, P(r) function using GNOM (Svergun, 1992). RAW uses MoW and Vc M.W. methods (Rambo & Tainer, 2013; Piiadov et al., 2018) |

**Table S3: BLI kinetic analysis of FakA-FakB2 interaction.**

1:1

| FakA Conc. (nM) | Response | KD (M) | KD Error | ka (1/Ms) | ka Error | kdis (1/s) | kdis Error | Full X^2 | Full R^2 |
|---|---|---|---|---|---|---|---|---|---|
| 1.65 | 0.0833 | 4.34E-09 | 4.58E-11 | 1.58E+05 | 1.16E+03 | 6.86E-04 | 5.21E-06 | 14.6797 | 0.9733 |
| 4.94 | 0.1297 | 4.34E-09 | 4.58E-11 | 1.58E+05 | 1.16E+03 | 6.86E-04 | 5.21E-06 | 14.6797 | 0.9733 |
| 14.8 | 0.1748 | 4.34E-09 | 4.58E-11 | 1.58E+05 | 1.16E+03 | 6.86E-04 | 5.21E-06 | 14.6797 | 0.9733 |
| 44.4 | 0.3551 | 4.34E-09 | 4.58E-11 | 1.58E+05 | 1.16E+03 | 6.86E-04 | 5.21E-06 | 14.6797 | 0.9733 |
| 133.3 | 0.6355 | 4.34E-09 | 4.58E-11 | 1.58E+05 | 1.16E+03 | 6.86E-04 | 5.21E-06 | 14.6797 | 0.9733 |

Bivalent Analysis (1:2)

| FakA Conc. (nM) | Response | KD (M) | KD Error | ka (1/Ms) | ka2 | ka Error | ka2 Error | kdis (1/s) | kdis2 | kdis Error | kdis2 Error | Full X^2 | Full R^2 |
|---|---|---|---|---|---|---|---|---|---|---|---|---|---|
| 1.65 | 0.0833 | 6.65E-09 | 5.47E-10 | 1.32E+05 | 9.36E+00 | 4.83E+03 | 7.68E-01 | 8.77E-04 | 5.06E-01 | 6.46E-05 | 7.57E-02 | 2.552 | 0.9768 |
| 4.94 | 0.1297 | 6.65E-09 | 5.47E-10 | 1.32E+05 | 9.36E+00 | 4.83E+03 | 7.68E-01 | 8.77E-04 | 5.06E-01 | 6.46E-05 | 7.57E-02 | 2.552 | 0.9768 |
| 14.8 | 0.1748 | 6.65E-09 | 5.47E-10 | 1.32E+05 | 9.36E+00 | 4.83E+03 | 7.68E-01 | 8.77E-04 | 5.06E-01 | 6.46E-05 | 7.57E-02 | 2.552 | 0.9768 |
| 44.4 | 0.3551 | 6.65E-09 | 5.47E-10 | 1.32E+05 | 9.36E+00 | 4.83E+03 | 7.68E-01 | 8.77E-04 | 5.06E-01 | 6.46E-05 | 7.57E-02 | 2.552 | 0.9768 |
| 133.3 | 0.6355 | 6.65E-09 | 5.47E-10 | 1.32E+05 | 9.36E+00 | 4.83E+03 | 7.68E-01 | 8.77E-04 | 5.06E-01 | 6.46E-05 | 7.57E-02 | 2.552 | 0.9768 |

Top: Octet Kinetics Table with 1:1 (1 FakB2:1 FakA binding site) binding model used for analysis. Bottom: Octet Kinetics Table with 1:2 (Bivalent Analyte, 1 FakB2: 2 FakA binding sites) binding model used for analysis.

**Table S4.** Select bacterial strains and plasmids used in this study.

| Strain or plasmid | Relevant characteristics[a, b] | Reference |
|---|---|---|
| **Strain name** | | |
| AH1263 | *S. aureus* wild-type USA300 | (32) |
| JLB2 | Δ*fakA* | (8) |
| JLB24 | *hla::Tn* | (8) |
| RN4220 | Restriction-deficient *S. aureus* strain | (33) |
| **Plasmids** | | |
| pCK2 | pCM28::*fakA*$^{D38A,\ D40A}$ | (10) |
| pCK10 | pET28a:: *fakA*$^{D38A,\ D40A}$ | This study |
| pCK13 | pCM28::*fakA*-His | (10) |
| pCK23 | pET28a::His-*fakB2* | This study |
| pCM28 | *E. coli*–*S. aureus* shuttle vector | (34) |
| pJB1036 | pCM28::*fakA*$^{C240A}$-His | This study |
| pJB1042 | pET28a::*fakA*$^{238\text{-}548}$ (c-term and middle) | This study |
| pJLB11 | pET28a::His-*fakA* | (9) |
| pJB165 | pCM28::*fakA* | (8) |
| pMJM2 | pET42a::GST-TEV-*fakA* | This study |
| pMJM3 | pET42a::GST-TEV-*fakA*$^{H282A,\ H284A}$ | This study |
| pMJM4 | pET42a::GST-TEV-*fakA*$^{C240A}$ | This study |
| pMJM5 | pET28a::TEV-His-*fakA*$^{328\text{-}548}$ (c-term) | This study |
| pMP4 | pCM28:: *fakA*$^{H282A,\ H284A}$-His | This study |

[a]pCM28 encodes chloramphenicol resistance

[b]pET vectors encode kanamycin resistance

[c]"Tn" Denotes an insertion mutation of the *bursa aurealis* transposon that is ErmR

**Table S5.** Oligonucleotides used in this study.

| Name | Sequence[a] | Reference |
|---|---|---|
| CNK24 | ggaattccatATGATTAGCAAAATTAATGGTAAATTATTTGC | This study |
| CNK25 | ccgctcgagTTATTCTACTGAAAAGAAATATTGATAAATTG | This study |
| CNK26 | ggctgcagTTAATGATGATGATGATGATGTTCTACTGAAAAGAAATATTG | (10) |
| CNK39 | ggaattccatATGACAAAACAGATTATAGTAACAGAC | This study |
| CNK40 | ccgctcgagTTACTTCTTAAGGACTACGAGG | This study |
| JBHEM1 | ccggatccATGAGTACCTCCTTTAATAAATATAAATACAC | (8) |
| JBKU81 | ATGGCTATGCTACTGAAATGATGGTTCG | This study |
| JBKU82 | CGAACCATCATTTCAGTAGCATAGCCAT | This study |
| JBKU104 | aggcatatgGGCTATTGTACTGAAATGATGGTTCG | This study |
| JB42 | ggctcgagTTATTCTACTGAAAAGAAATATTGATAAATTG | (8) |
| MP7 | GTGAAAGTTGCTGTGGCTACCGAATAC | This study |
| MP11 | GTATTCGGTAGCCACAGCAACTTTCACAATTTCTTCATC | This study |

[a]Sequences are provided 5’ to 3’ and lower-case letters denote bases added for cloning purposes

**Table S6.** NanoDrop One UV-Vis Spectrophotometer Parameters

| **Protein Construct:** | **MW (Da):** | **Theoretical pI** | **ε ($M^{-1}$ $cm^{-1}$)** |
|---|---|---|---|
| His-FakA | 62678.99 | 4.86 | 26610 |
| His-FakB2 | 32808.21 | 6.08 | 13410 |
| GST-TEV-FakA | 87382.73 | 4.87 | 71210 |
| Untagged FakA | 60515.67 | 4.67 | 26610 |
| His-FakA$^{328\text{-}548}$ (c-term) | 26450.68 | 4.44 | 14440 |
| His-FakA$^{238\text{-}548}$ (middle and c-term) | 37193.03 | 4.95 | 19035 |

**Table S7**. Crystallographic data for the FakA N-terminal domain structures.

| Structure<br>PDB Code | Mn-FakA_N<br>8VIP | AMP-PNP-FakA_N<br>8VIQ | Apo-FakA_N<br>8VIR | ADP-FakA_N<br>8VIT |
|---|---|---|---|---|
| **Data Collection** | | | | |
| Unit-cell parameters (Å, °) | *a*=42.81<br>*b*=57.03<br>*c*=81.70 | *a*=42.73<br>*b*=56.43<br>*c*=81.80 | *a*=42.05<br>*b*=47.54<br>*c*=93.23 | *a*=42.94<br>*b*=57.09<br>*c*=81.03 |
| Space group | $P2_12_12_1$ | $P2_12_12_1$ | $P2_12_12_1$ | $P2_12_12_1$ |
| Resolution (Å)[1] | 46.76-1.77 (1.81-1.77) | 46.45-1.10 (1.12-1.10) | 47.54-1.90 (1.94-1.90) | 46.67-1.25 (1.27-1.25) |
| Wavelength (Å) | 1.7712 | 1.0000 | 1.0000 | 1.0000 |
| Temperature (K) | 100 | 100 | 100 | 100 |
| Observed reflections | 206,922 | 512,550 | 122,554 | 353,128 |
| Unique reflections | 19,656 | 81,027 | 15,360 | 55,921 |
| $\langle I/\sigma(I)\rangle$[1] | 24.2 (2.5) | 12.8 (1.6) | 11.0 (1.9) | 14.2 (1.5) |
| Completeness (%)[1] | 97.2 (72.5) | 100 (100) | 100 (100) | 100 (100) |
| Multiplicity[1] | 10.6 (2.6) | 6.3 (6.2) | 8.0 (8.2) | 6.3 (6.0) |
| $R_{merge}$ (%)[1, 2] | 6.5 (33.5) | 5.6 (113.3) | 12.0 (112.9) | 5.3 (115.2) |
| $R_{meas}$ (%)[1, 4] | 6.8 (40.4) | 6.1 (123.7) | 12.9 (120.6) | 5.8 (126.3) |
| $R_{pim}$ (%)[1, 4] | 2.0 (22.0) | 2.4 (49.2) | 4.6 (42.1) | 2.3 (51.1) |
| $CC_{1/2}$[1, 5] | 0.999 (0.884) | 0.999 (0.675) | 0.998 (0.664) | 0.999 (0.754) |
| DelAnom CC[6] | 0.359 | - | - | - |
| **Refinement** | | | | |
| Resolution (Å)[1] | 46.76-1.77 | 34.07-1.10 | 42.35-1.90 | 34.32-1.25 |
| Reflections (working/test)[1, 7] | 34,454/1,756 | 76,964/3,958 | 14,584/726 | 53,071/2,753 |
| $R_{factor}$ / $R_{free}$ (%)[1,3] | 14.8/18.6 | 12.5/14.2 | 17.4/22.0 | 13.8/16.1 |
| No. of atoms (Protein/AMP-PNP/Mg or Mn/Water) | 1,536/31/2/143 | 1,596/31/2/229 | 1,526/-/-/85 | 1,585/27/2/18<br>6 |
| **Model Quality** | | | | |
| R.m.s deviations | | | | |
| Bond lengths (Å) | 0.009 | 0.006 | 0.010 | 0.006 |
| Bond angles (°) | 1.092 | 0.988 | 0.921 | 0.980 |
| Mean *B*-factor (Å$^2$) | | | | |
| All Atoms | 22.6 | 17.8 | 28.5 | 20.5 |
| Protein | 21.9 | 15.5 | 28.2 | 19.2 |
| AMP-PNP | 21.1 | 12.8 | - | 14.5 |
| Mg or Mn | 20.2 | 16.7 | - | 16.3 |
| Water | 31.4 | 34.3 | 33.7 | 31.6 |
| Coordinate error (maximum likelihood) (Å) | 0.16 | 0.09 | 0.26 | 0.11 |
| Ramachandran Plot | | | | |
| Most favored (%) | 98.6 | 98.6 | 98.6 | 98.1 |
| Additionally allowed (%) | 1.4 | 1.4 | 1.4 | 1.9 |

1) Values in parenthesis are for the highest resolution shell.
2) $R_{merge} = \Sigma_{hkl}\Sigma_i |I_i(hkl) - \langle I(hkl)\rangle| / \Sigma_{hkl}\Sigma_i I_i(hkl)$, where $I_i(hkl)$ is the intensity measured for the *i*th reflection and $\langle I(hkl)\rangle$ is the average intensity of all reflections with indices hkl.
3) $R_{factor} = \Sigma_{hkl} ||F_{obs}(hkl)| - |F_{calc}(hkl)|| / \Sigma_{hkl} |F_{obs}(hkl)|$; Rfree is calculated in an identical manner using 5% of randomly selected reflections that were not included in the refinement.
4) $R_{meas}$ = redundancy-independent (multiplicity-weighted) $R_{merge}$[1, 2]. $R_{pim}$ = precision-indicating (multiplicity-weighted) $R_{merge}$[3, 4].
5) $CC_{1/2}$ is the correlation coefficient of the mean intensities between two random half-sets of data [5, 6].
6) DelAnom CC is the correlation coefficient between the Bijvoet differences ($I_{(hkl)} - I_{(-h-k-l)}$) from two random half-sets of data [1] and is used to estimate the anomalous signal strength.
7) For the FakA-Mn structure, the number of reflections used for refinement (34,454) is greater than the number of unique reflections obtained after scaling the data (19,656). This results from the treatment of Friedel pairs (F+/F-) as unique reflections during refinement in order to refine the anomalous scattering factors for the $Mn^{2+}$ ions.

**Expression and purification of FakA, FakA variants, and FakB2**. Each *S. aureus* protein was overexpressed in BL21 (DE3) *Escherichia coli* in 2X-YT media in baffled Fernback flasks, shaking at 200 RPM or in a Lex-48 bioreactor at 37°C*.* Induction of expression was started after an OD600 of ~0.6 (about 4 hours) was reached with 1 mM isopropyl-β-D-thiogalactopyranoside (IPTG). Cells were harvested after 4 additional hours at 37°C and frozen at -20°C. For His-tagged constructs, the bacterial pellet was resuspended in a buffer containing 50 mM Tris [pH 7.4], 50 mM KCl, and 5 mM Imidazole with a mixture of protease inhibitors (Leupeptin, AEBSF, Pepstatin A, Benzamidine and 1 mM β-Mercaptoethanol). This suspension was sonicated on ice and centrifuged at 24,424xg for 1 hour to collect cell lysate. The lysate was filtered through a 0.45-micron filter and loaded onto a HisTrap HP 5 mL column. This column was equilibrated with Buffer HisA (50 mM Tris [pH 7.4], 50 mM KCl, 5 mM Imidazole) and bound proteins were eluted with Buffer HisB (50 mM Tris [pH 7.4], 50 mM KCl, 500 mM Imidazole). Peak fractions containing our target protein were combined and diluted using Buffer HisA and run through a Hitrap Q HP column with Buffer QA (50 mM Tris [pH 7.4], 50 mM KCl, 10% glycerol) and eluted with Buffer QB (50 mM Tris [pH 7.4], 1 M KCl, 10% glycerol). The fractions containing the purest FakA protein, as determined by SDS-PAGE, were run on a HiPrep 16/60 Sephacryl S-200 HR gel filtration column with 50 mM Tris [pH 7.4], 150 mM KCl, 1 mM TCEP, and 5% glycerol. The fractions containing the purest FakA were again determined by SDS-PAGE, concentrated using a 10 or 30K cutoff, flash frozen with liquid nitrogen, and stored at -80°C. Final concentrations were determined via A280 using a NanoDrop One UV-Vis Spectrophotometer (Table S6).

For GST-TEV-tag constructs*,* the bacterial pellet was resuspended in 50 mM Tris [pH 7.4], 200 mM KCl, and 1 mM β-Mercaptoethanol. Resuspended cells were treated with the same protease inhibitors and disrupted to produce cell lysates as described above. The target protein was first purified using a GSTPrep FF 16/10 column with Buffer GSTA (50 mM Tris [pH 7.4], 200 mM KCl, 1 mM β-Mercaptoethanol) and eluted with Buffer GSTB (50 mM Tris [pH 7.4], 200 mM

KCl, 1 mM β-Mercaptoethanol, 50 mM glutathione). Fractions containing GST-FakA as determined by SDS-PAGE were combined and cut to remove the GST-tag with Tobacco Etch Virus (TEV) protease at room temperature overnight. TEV cutting was confirmed through SDS-PAGE and protein was buffer exchanged/concentrated into Buffer GSTA. Protein was separated using a GSTPrep FF 16/10 column, this time collecting the flow-through (cut FakA) and running peak fractions on SDS-PAGE to determine untagged FakA-containing fractions. Gel filtration using a HiPrep 16/60 Sephacryl S-200 HR column as described above was performed on these FakA-containing fractions, the protein was concentrated using a 30K cutoff, and stored as described above.

**Limited proteolysis:** X-ray crystallography was incompatible with full-length FakA purified protein. Thus, limited proteolysis was performed to identify organized domains to be the N-terminal domain (Met1-Ala212), Middle domain (Lys213-Lys327), and C-terminal domain (Met328-Glu548).

**X-ray Crystallography.** A purified construct of the FakA N-terminal domain (FakA-N, M1-A212) was concentrated to 10.1 mg $mL^{-1}$ (0.4 mM) in 150 mM KCl, 50 mM Tris [pH 7.4], 1 mM TCEP for crystallization screening. All crystallization experiments were set up using an NT8 drop setting robot (Formulatrix Inc.) and UVXPO MRC (Molecular Dimensions) sitting drop vapor diffusion plates at 18°C using 100 nL of protein and 100 nL of well solution equilibrated against 50 μL of the latter. Samples in complex with ligands were prepared by adding 2.5 mM $MgCl_2$ or $MnCl_2$ and 2.5 mM ADP or AMP-PNP to the protein and incubating on ice for 30 minutes before screening. Prismatic crystals used for X-ray diffraction data collection were obtained as follows. **Apo-FakA_N**: Proplex screen (Molecular Dimensions) condition D1 (25% (w/v) PEG 4000, 100 mM HEPES [pH 7.5], 200 mM NaCl). **ADP-FakA_N:** Proplex screen (Molecular Dimensions) condition C3 (20% (w/v) PEG 4000, 100 mM sodium acetate [pH 5.0], 200 mM ammonium acetate). Samples were cryoprotected in a solution composed of 80% well solution and 20% (v/v) PEG 200 which was dispensed onto the drop, crystals were harvested immediately and stored in liquid nitrogen. **AMP-PNP-FakA_N and Mn-FakA_N:** Proplex screen (Molecular Dimensions) condition D9 (15% (w/v) PEG 6000, 100 mM MES [pH 6.5], 5% (v/v) 2-methyl-2,4-pentanediol (MPD)). Crystals of **Mn-FakA_N** were obtained in an identical manner as described above except that 2.5 mM $MnCl_2$ and 2.5 mM AMP-PNP were added to an aliquot of the protein and crystals were obtained from Proplex D9 in CombiClover 300 sitting drop vapor diffusion plates (Rigaku Reagents). A cryoprotectant solution composed of 85% crystallization solution and 15% (v/v) MPD was dispensed onto the drops, and crystals were harvested immediately and frozen. X-ray diffraction data were collected at the Advanced Photon Source beamline 17-ID using a Dectris Pilatus 6M pixel array detector.

Intensities were integrated using XDS [1, 2] via Autoproc [3] and the Laue class analysis and data scaling were performed with Aimless [4]. Structure solution was conducted by SAD phasing with Crank2 [5] using the Shelx [6], Refmac [7], Solomon [8], Parrot [9], and Buccaneer [10] pipeline via the CCP4 [11] interface using a Mn-FakA_N data set. Two highly occupied $Mn^{2+}$ ions were located that yield peaks in the anomalous difference map at $60\sigma$ and $38\sigma$ levels. Additionally, contributions to the anomalous signal were obtained from the sulfur atoms of six methionine residues, two cysteine residues, and two phosphate atoms from AMP-PNP, which produced peaks between $5\sigma$ to $10\sigma$ in the anomalous difference map. Phasing/density modification resulted in a mean figure of merit of 0.76. Subsequent model building with density modification and phased refinement yielded $R/R_{free}$ = 0.24/0.27 for the refined model (212 residues). Additional refinement and manual model building were conducted with Phenix [12] and Coot [13], respectively. The Mn-FakA_N model was used for molecular replacement with Phaser [14] against the higher resolution AMP-PNP-FakA_N data set and the top solution was obtained in the space group $P2_12_12_1$ (TFZ=78.4, LLG=16,757 AMP-PNP-FakA_N). The model was further improved by automated model building Phenix and Coot. Anisotropic atomic displacement parameters were refined for all atoms. The final model was used for molecular replacement searches against the Apo-FakA_N and ADP-FakA_N data sets. Disordered side chains were truncated to the point for which electron density could be observed. Structure validation was conducted with Molprobity [15] and figures were prepared using the CCP4MG package [16]. Structure superposition was carried out using Gesamt [17]. Crystallographic data are provided in Table S7.

**Size exclusion chromatography coupled to Small Angle X-ray Scattering (SEC-SAXS).** Small angle X-ray scattering was performed at BioCAT (beamline 18ID-D at the Advanced Photon Source, Argonne National Laboratory) with in-line size exclusion chromatography (SEC-SAXS) to separate sample from aggregates and other contaminants thus ensuring optimal sample quality. The sample was loaded onto a Superdex 200 Increase 10/300 GL column (Cytiva), which was run at 0.6 ml $min^{-1}$ by an AKTA Pure FPLC (GE), and the eluate after it passed through the UV monitor was flown through the SAXS flow cell. The flow cell consists of a 1.0 mm ID quartz capillary with ~20 μm walls. A coflowing buffer sheath is used to separate samples from the capillary walls, helping prevent radiation damage [18]. Scattering data was recorded using a Pilatus3 X 1M (Dectris) detector which was placed 3.663 m from the sample giving us access to a q-range of 0.0029 $Å^{-1}$ to 0.42 $Å^{-1}$ ($q=4\pi\sin\theta/\lambda$, where $\lambda$ is the wavelength and $2\theta$ is the scattering angle). 0.5 second exposures were acquired every 1 second during elution and data was reduced using BioXTAS RAW 2.1.4 [19]. Buffer blanks were created by averaging regions before the elution peak and subtracted from exposures selected from the elution peak, which were deconvoluted using evolving factor analysis (EFA) with default settings as implemented in BioXTAS RAW 2.1.4 [20] to create the I(q) vs q curves used for subsequent analyses. The validity of the overall deconvolution was assessed on the component concentration profiles and mean error-weighted $\chi^2$ for the whole deconvolution range.

The forward scattering intensity I(0), and the radius of gyration ($R_g$) were calculated from the Guinier fit. The normalized Kratky plot, the pair-distance distribution plot P(r), and the corrected Porod volume were calculated using GNOM embedded in BioXTAS RAW 2.1.4 [21]. Low-resolution ab initio bead modeling was carried out using 20 reconstructions by DAMMIF [22] in either P1 or P2 symmetry with fast or slow mode, averaged by DAMAVER [23], and a final structure was refined in DAMMIN[24]. AMBIMETER was used to assess the ambiguity of reconstructions [22]. Electron density reconstructions were performed using DENSS program [25], as implemented in BioXTAS RAW 2.1.4. The DENSS densities have a Fourier shell correlation (FSC) of 49.6 Å. Molecular dynamics (MD) based modeling was performed using BilboMD [26] implemented in the web server (https://bl1231.als.lbl.gov/bilbomd) using 400 conformations per $R_g$ value, $R_g$ range 37-47 Å from experimental $R_g$ determined from $P(r)$ function, flexible fragments defined manually. Docking of generated models with the SAXS-derived bead-based modeling or electron density modeling was done manually and improved by rigid-body refinement using Chimera [27]. The calculation of theoretical scattering curves for the models was performed by the program CRYSOL [28], which also determines the discrepancy ($\chi^2$ value) between the simulated and experimental scattering curves. SEC-SAXS data collection and analysis statistics are in Tables S1 and S2.
